# Powdery mildew fungi block plant vacuolar traffic to suppress immunity

**DOI:** 10.64898/2026.05.12.724530

**Authors:** Sohini Deb, João P. A. Plácido, Xuan Li, Paraskevi Doukoudaki, Björn Sabelleck, Valeria Velásquez-Zapata, Gregory Fuerst, Bobby Routya, J. Mitch Elmore, Anna Christensen, Roger P. Wise, Hans Thordal-Christensen

## Abstract

Plant immunity can be activated by membrane-localized pattern recognition receptors or by cytosolic nucleotide-binding leucine-rich repeat (NLR) receptors, while it can be counteracted by pathogen secreted effectors. Manifestation of immunity often involves endomembrane traffic. However, limited evidence is available for such an involvement in NLR-mediated immunity, which typically includes a hypersensitive reaction (HR)-programmed cell death response. Here we show that the barley powdery mildew fungus uses several effectors to target and inhibit the vacuolar trafficking pathway, which causes endomembrane markers to stall in the endoplasmic reticulum (ER). We used this ER-stalling phenotype as a proxy to track fungal manipulation of the vacuolar pathway during different stages of the actual infection. Our data indicate that powdery mildew fungi interfere with the vacuolar pathway, but only temporarily, as the stalling is lifted once the fungal haustorial feeding structures are well-developed for nutrient uptake in epidermal cells. Notably, we show evidence that blocking the vacuolar pathway causes a general inhibition of NLR-mediated HR, and that this mechanism is taken advantage of by the pathogen. Finally, we provide an example that the fungus can secrete a different set of effectors at the haustorial stage that result in the re-opening of the vacuolar pathway by inhibiting the initial vacuolar traffic-suppressing effectors.

## INTRODUCTION

Powdery mildew fungi are obligate biotrophic pathogens that colonize a wide range of angiosperm species (Braun and Cook, 2012). An example is the model pathogen, *Blumeria hordei* (*Bh*), specialized to cause disease in barley (*Hordeum vulgare*). *Bh* attacks by forming an appressorium, which serves to penetrate the epidermal cell of the host, followed by the formation of a feeding structure, termed haustorium, inside the host cell (Hückelhoven and Panstruga, 2011). Like other pathogens, powdery mildew fungi transfer large numbers of effector proteins into the plant cell to target and inhibit immune components and manipulate the cell to accommodate the haustorium and feed the fungus (Yuen et al., 2023).

Plants recognize pathogen attacks through plasma membrane (PM)-localized pattern recognition receptors or cytosolic nucleotide-binding leucine-rich repeat (NLR) receptors, which initiate the two distinct, but highly connected, responses: pattern-triggered immunity (PTI) and effector-triggered immunity (ETI), respectively (Jones et al., 2024). NLR receptors, of either the ‘Toll-interleukin receptor (TIR)-NLR’ (TNL) or the ‘Coiled-coil (CC)-NLR’ (CNL) type, detect effectors directly or indirectly and activate ETI, often resulting in a programmed cell death response termed the ‘hypersensitive reaction’ (HR). Activated CNLs, such as Arabidopsis ZAR1 and wheat Sr35, form ‘resistosomes’, Ca^2+^-permeable pores that are inserted into the PM (Bi et al., 2021; Förderer et al., 2022). Activated TNLs, on the other hand, form resistosomes with enzymatic activity whereby nucleotide-based signaling molecules are synthesized, triggering EDS1/PAD4 and EDS1/SAG101 heterodimers to interact with the helper RPW8-type NLRs (RNLs) of the ADR1 and NRG1 subclasses, respectively. This, in turn, triggers the RNLs to form Ca^2+^-conducting pores as well (Locci and Parker, 2024). The processes between the initial cytosolic Ca^2+^-influx and HR/pathogen resistance during ETI are still poorly understood.

The early steps of the endomembrane trafficking pathway in plants usually involve transport of cargo proteins through the ER and Golgi, where proteins undergo several maturation steps. The *trans*-Golgi network (TGN) serves as a subsequent sorting hub, where proteins can be sorted for secretion or towards the vacuole (Shimizu et al., 2021). The canonical vacuolar pathway in plants involves the formation of a Rab5 GTPase-labeled multivesicular body (MVB), which later matures into a Rab7 GTPase-labeled MVB (Singh et al., 2014). Throughout this process, the endosomal sorting complex required for transport (ESCRT) machinery generates MVB intraluminal vesicles (ILVs), carrying cargo proteins destined for vacuolar degradation (Haas et al., 2007). In plants, unlike other systems, the class C core vacuole/endosome tethering (CORVET) complex is thought to mediate fusions between the Rab5-positive MVB and the tonoplast, while the homotypic fusion and vacuole protein sorting (HOPS) complex is thought to mediate fusion between Rab7-positive MVBs and the tonoplast (Takemoto et al., 2018).

We have previously suggested ETI and vacuolar traffic to be linked. This was based on studies of the effector *Bh*CSEP0162 and its target, MON1, which is the Rab7 guanine nucleotide exchange factor (GEF), and the core effector *Bh*CSEP0214 and its target, VPS18, which is a shared component of CORVET and HOPS. The studies suggested that ETI against *Bh*, mediated by allelic MLA CNLs in barley, is dependent on the vacuolar pathway being functional (Liao et al., 2023; Sabelleck et al., 2025). In addition, we have found that CNL-mediated immunity in Arabidopsis and *Nicotiana benthamiana* requires a functional ESCRT-III machinery (Schultz-Larsen et al., 2018). However, the extent by which powdery mildew fungi manipulate this trafficking pathway during infection has remained unexplored. Here, we demonstrate that *Bh* uses multiple effectors to temporarily block vacuolar traffic, as observed by effective stalling of proteins in the ER, and thereby disrupts NLR-mediated immunity.

## RESULTS

### Inhibition of the pathway to the vacuole stalls its markers in the ER

We have previously shown that overexpression of CSEP0214 and silencing of its target, VPS18, cause ER-stalling of the vacuolar cargo marker, (SP)-RFP-AFVY (Scheuring et al., 2011), in epidermal cells of the monocot plant, barley (Sabelleck et al., 2025). Here we first asked whether such an ‘ER-stalling’ is a common response to the arrest of the vacuolar pathway. Thus, we used the dominant-negative mutant version of the ESCRT-III VPS4 AAA-ATPase, VPS4^E232Q^ (Haas et al., 2007; Pfitzner et al., 2020) (hereon referred to as VPS4^EQ^) as a tool to inhibit vacuolar traffic. VPS4 sequentially disassembles ESCRT-III to allow its subunit exchange during ILV formation in MVBs (Pfitzner et al., 2020). We observed that VPS4^EQ^ also leads to the mis-localization of (SP)-RFP-AFVY to reticular and perinuclear structures in barley, where it overlaps with ERD2-mYFP, an ER/ER-exit-site/Golgi marker derived from the HDEL ER-retention signal receptor (Hanton et al., 2007; 2008) (Figure 1A). To confirm this consequence of VPS4^EQ^ expression, we additionally analyzed another marker in the same pathway, the broadly used Golgi marker, ST-YFP (Brandizzi et al., 2002; Sabelleck et al., 2025), and saw that it was also stalled in the ER after VPS4^EQ^ expression, indicating that the first ER-to-Golgi step in the pathway is affected (Figure 1B), despite it being several pathway steps earlier to where VPS4 functions. This is reminiscent of how dominant-negative Sar1 GTPase causes Golgi structures to collapse into the ER (Ito et al., 2012). To analyze this phenomenon in a dicot plant, we expressed these proteins in leaf epidermal cells of *N. benthamiana* after *Agrobacterium*-mediated transformation. Here VPS4^EQ^ caused (SP)-RFP-AFVY to be partly secreted, as described in similar studies (Hu et al., 2022; Singh et al., 2014), and to be partly stalled in an ER-like reticular structure, comparable to the phenotype in barley (Figure 1C). VPS4^EQ^ also caused ST-YFP to be stalled in the ER, overlapping with (SP)-mCherry-HDEL, in *N. benthamiana* (Figure 1D). As an example from outside the plant kingdom, we studied a strain of yeast (*Saccharomyces cerevisiae*) mutated in the *Vps18* gene. Interestingly, (SP)-RFP-AFVY was vacuolar-localized in wild-type yeast and ER-localized in the *vps18* mutant (Figure 1E). We further tested another vacuolar marker, (SP)-RFP-QRPL (Casler and Glick, 2020), and found that the ER-stalling effect in the yeast *vps18* mutant is not limited to the AFVY vacuolar-localization signal (Supplemental Figure 1). Taken together, our results demonstrate that the block of protein traffic through the vacuolar pathway results in a build-up already at the initial sorting step in the ER. Importantly, the ER-stalling phenotype appears to be highly conserved, and therefore it is a useful proxy for studying how pathogens manipulate vacuolar traffic.

**Figure 1.**
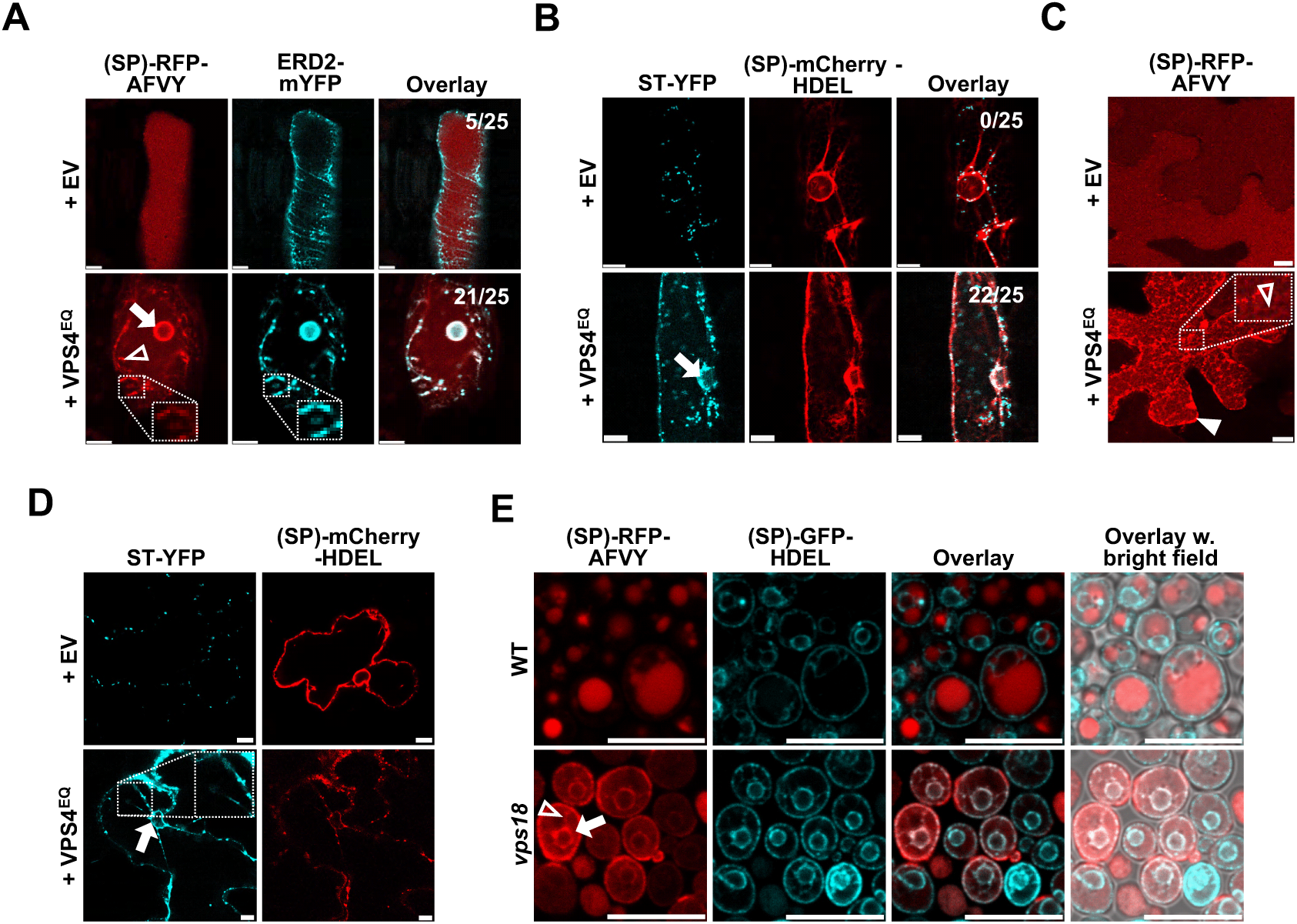
Markers of the vacuolar transport pathway are stalled in the ER when the endomembrane pathway is inhibited. **(A and B)** Vacuolar marker [(SP)-RFP-AFVY] and Golgi marker (ST-YFP) stalled in ER, with reticular (insert) and perinuclear signal overlapping with the ER markers, ERD2-mYFP and (SP)-mCherry-HDEL, respectively, when co-expressed with dominant-negative ESCRT-III component, VPS4^E232Q^, in barley *cv.* Golden Promise leaf epidermal cells by particle bombardment. Numbers indicate proportions of cells showing ER stalling. **(C and D)** Vacuolar marker [(SP)-RFP-AFVY] and Golgi marker (ST-YFP) stalled in ER with reticular (insert), perinuclear and extracellular signal in *N. benthamiana* leaf epidermal cells when co-expressed with VPS4^E232Q^ by *Agrobacterium* infiltration of 3-week-old *N. benthamiana* plants. Cells were imaged 2 d after infiltration. **(E)** Vacuolar marker [(SP)-RFP-AFVY] stalled in ER with reticular and perinuclear signal overlapping with the ER marker [(SP)-GFP-HDEL] in stably transformed *vps18* mutant, but not in WT, of yeast strain BY4741. EV, empty vector. Open arrowhead, reticular signal overlapping with ER marker. Closed arrow, perinuclear signal overlapping with the ER marker. Closed arrowhead, extracellular signal. Scale bars, 10 μm. These results are representative of the outcomes of at least three independent experiments.

### Barley powdery mildew effectors targeting the vacuolar pathway cause ER-stalling of vacuolar markers

Having established that the *Bh*CSEP0162 and *Bh*CSEP0214 effectors target MON1 and VPS18, respectively, in the vacuolar trafficking pathway (Liao et al., 2023; Sabelleck et al., 2025), we expanded our search for more of such *Bh* effector targets with our yeast two-hybrid next-generation interaction screening pipeline (Y2H-NGIS) (Elmore et al., 2023; Li et al., 2024; Smith et al., 2025; Velásquez-Zapata et al., 2021; 2023). Here we found highly significant Y2H-SCORES between *Bh*CSEP0174 and the barley Adaptor Protein1 gamma2 (AP1γ2), and between *Bh*CSEP0219 and five different Seven IN Absentia (SINA) E3 ligases (Supplemental Data Table 1). These results were subsequently verified by binary Y2H and *in planta* bi-fluorescence complementation (Supplemental Figure 2).

Both AP1γ2 and SINA E3 ligases function in the vacuolar trafficking pathway. AP1γ2 is involved in vesicle formation at the TGN (Wang et al., 2014), while SINA E3 ligases ubiquitinate the ESCRT-I proteins, FREE1 and VPS23a, required for ESCRT function (Gao et al., 2017; Xia et al., 2020). Similar to *Bh*CSEP0214 (Sabelleck et al., 2025), we observed that overexpression of *Bh*CSEP0162, *Bh*CSEP0174 or *Bh*CSEP0219 led to ER-stalling of (SP)-RFP-AFVY in barley epidermal cells, as confirmed by co-localization with ERD2-mYFP (Figure 2). To a lesser extent, it also caused this vacuolar marker to be secreted to the extracellular space (Supplemental Figure 3A). Moreover, these four effectors also caused stalling of the Golgi marker, ST-YFP, in the ER of barley cells (Supplemental Figure 3B), and ER-stalling of both (SP)-RFP-AFVY and ST-YFP in *N. benthamiana*, where secretion of the vacuolar marker was more pronounced (Supplemental Figures 4A and 4B). *Bh*CSEP0055, which interacts with plant extracellular pathogenesis-related proteins (Zhang et al., 2012), was used as negative control in these experiments.

**Figure 2.**
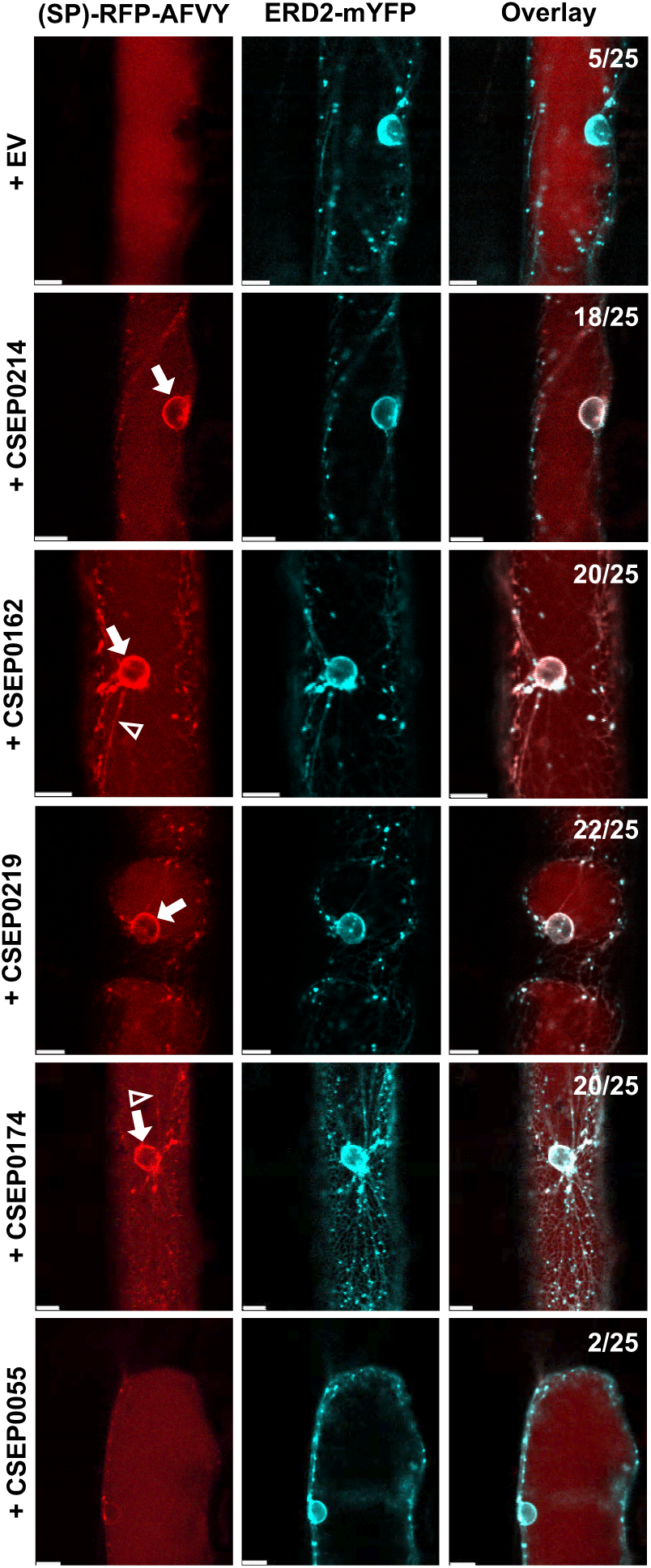
Barley powdery mildew effectors targeting the endomembrane trafficking pathway cause ER-stalling of vacuolar pathway marker. Vacuolar marker [(SP)-RFP-AFVY] stalled in ER with reticular and perinuclear signal overlapping with the ER marker, ERD2-mYFP, in barley *cv.* Golden Promise leaf epidermal cells when co-expressed with *Bh* effectors. CSEP0055, negative control effector not targeting endomembrane traffic. EV, empty vector. Numbers indicate proportions of cells showing ER stalling. Cells were imaged 2 d after transient expression by particle bombardment. Open arrowhead, reticular signal overlapping with ER marker. Closed arrow, perinuclear signal overlapping with the ER marker. Scale bars, 10 μm. These results are representative of the outcomes of at least three independent experiments.

In agreement with the ER-stalling caused by these effectors, we observed the same consequence of silencing their host targets, *MON1*, *AP1γ2*, *SINA6* and *VPS18* (Supplemental Figure 5), as shown previously for the latter (Sabelleck et al., 2025). The reduced ER-stalling rate observed after the silencing of *SINA6* was likely due to redundancy caused by other SINA E3 ligases. Taken together, this suggests that these powdery mildew effectors block the vacuolar marker in the ER by targeting distinct steps of the vacuolar pathway.

### Blocking vacuolar traffic indirectly inhibits NLR-mediated HR

*Pseudomonas* type III secretion-mediated delivery of *Bh*CSEP0214 impedes barley CNL MLA1 and MLA3-activated HR during *Bh* attack (Sabelleck et al., 2025). Here we further tested *Bh*CSEP0162, *Bh*CSEP0174 and *Bh*CSEP0219 and found that they also suppress MLA3-mediated cell death and enhance fungal growth in the P02 (*Mla3*) / *Bh*A6 (*AVR_a3_*) incompatible interaction (Figures 3A and 3B). Considering that all these effectors also appear to inhibit vacuolar traffic, the results suggest that blocking this pathway affects MLA-mediated HR. This was further confirmed in *N. benthamiana*, where HR induced by the MLA1-AVR_A1_ CNL-effector pair was suppressed by expression of VPS4^EQ^ (Figures 3C and 3D). To test this for other NLRs, we also included the widely analyzed Arabidopsis CNL, RPM1 (Gao et al., 2011). Two ESCRT-III components, VPS4 and the AMSH3 de-ubiquitinase, are required for HR in *N. benthamiana* induced by the constitutively active RPM1^D505V^ (RPM1^DV^) (Schultz-Larsen et al., 2018). Here we used the same setup to quantitively assay HR and confirmed that RPM1^DV^-activated HR can be suppressed by expression of VPS4^EQ^ (Figures 3E and 3F). Furthermore, treatment with Wortmannin, a well-characterized chemical suppressor of MVB formation (Ebine et al., 2011; Emans et al., 2002), also showed that RPM1^DV^-activated HR is inhibited when vacuolar traffic is blocked (Figures 3G and 3H). It is noteworthy that protein levels of RPM1 and RPM1^DV^, as well as their subcellular localization at the PM, remained unchanged upon expression of VPS4^EQ^ (Supplemental Figure 6).

**Figure 3.**
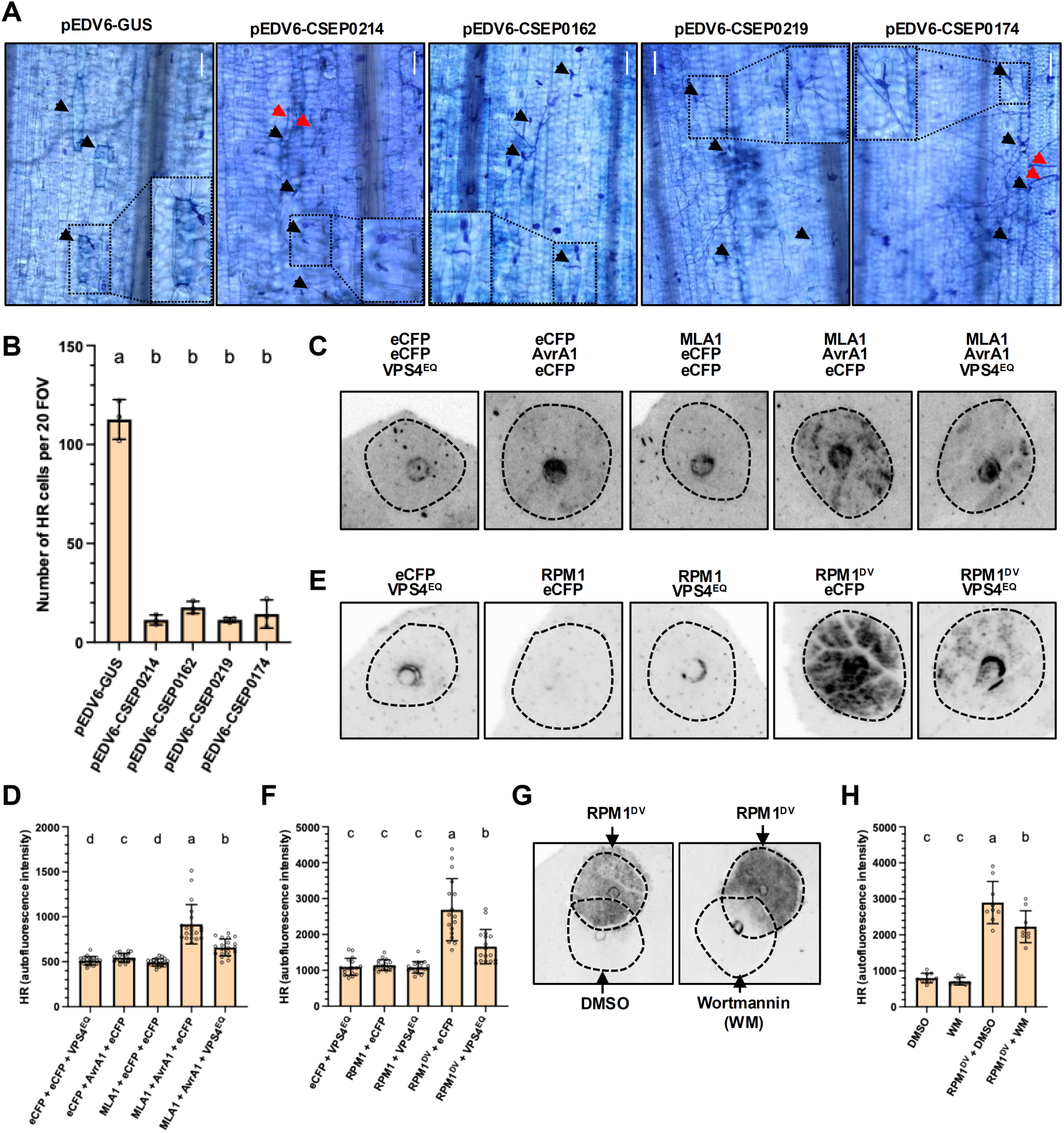
MLA3 and RPM1-mediated HR require endosomal trafficking. **(A and B)** CSEPs targeting plant endomembrane trafficking components inhibit MLA3-mediated resistance and HR against the *Bh* isolate A6 in the barley line P02. EtHAn bacteria for T3S-based introduction of GUS (negative control) and CSEPs were infiltrated immediately prior to inoculation with *Bh*. **(A)** Growth of fungal structures and HR cells visualized at 72 hai by staining with Trypan Blue and Coomassie Blue, respectively. Black arrowheads, attacked cells; red arrowheads, conidiophores. Inserts zoom in on single attacked cells. Scale bars, 100 µm. **(B)** HR cell death was quantified by counting the total number of *Bh*-attacked cells stained with Trypan Blue in 20 fields of view (FOV) per leaf sample (n=3). **(C, E and G)** Representative images of VPS4^EQ^-inhibition of MLA1-AvrA1 **(C)**, of RPM1^DV^ **(E)** and of Wortmannin **(G)**-inhibition of RPM1^DV^-mediated HR in *N. benthamiana*, visualized by red-light imaging 2 d after *Agrobacterium* infiltration and 1 d after estradiol-mediated induction of MLA and RPM1 expression. **(D, F and H)** Quantification by red-light imaging of HR in **(C)**, **(E)** and **(G)**. Average fluorescence intensity in the infiltrated areas was quantified using ImageJ. Error bars, ± SD; n=20 **(D)**; n=17 **(F)**; n=9 **(H)**. Wortmannin (30 μM) or DMSO were infiltrated 1 h after *Agrobacterium* infiltration. Dots represent individual data points. Different letters represent statistical differences between treatments assessed by unpaired **(B)** or paired **(D, F and H)** one-way ANOVA with Tukey’s honestly significant difference (HSD) (*P* < 0.05). These results are representative of the outcomes of at least three independent experiments.

Next we tested the requirement of this pathway for several other Arabidopsis NLRs. This revealed that HR activated by the CNL RPS2 (Mackey et al., 2003), the constitutively active CNL RPS5^D266E^ (Ade et al., 2007), the CC-domain of the CNL ZAR1 (Baudin et al., 2017), and by the RNL ADR1 (Saile et al., 2021), was inhibited by the expression of VPS4^EQ^ for 24 h prior to the expression of the NLR. In contrast, HR activated by the TIR domains of TNLs SNC1 and RPS4 (Bernoux et al., 2023), and by the constitutively active RNL NRG1.1^D485V^ (Jacob et al., 2021), was not inhibited under these conditions (Figures 4A and 4B). Surprisingly, extending the expression of VPS4^EQ^ to 48 h prior to NLR expression led to inhibition of HR mediated by all NLRs tested (Figure 4C and 4D). Blocking vacuolar traffic for 48 h by expressing the *Bh* effectors *Bh*CSEP0162, *Bh*CSEP0174, *Bh*CSEP0214, and *Bh*CSEP0219, as well as dominant-negative RING-domains of the HOPS components VPS18 and VPS41 (Sabelleck et al., 2025), all had a similar effect compared to VPS4^EQ^, namely inhibiting both RPM1^DV^- and SNC1^TIR^-mediated HR in *N. benthamiana* (Supplemental Figure 7). These results indicated that inhibition of HR by blocking vacuolar traffic is quantitative. We thus tested how blocking vacuolar traffic for even shorter time periods, using Wortmannin, affected RPM1^DV^-activated HR, and found that 1 h of pre-treatment did not inhibit HR, unlike the 24 h pre-treatment (Figures 4E and 4F). This is particularly surprising since a 30-min treatment with Wortmannin is usually sufficient for blocking the vacuolar pathway in plants (Emans *et al*., 2002).

**Figure 4.**
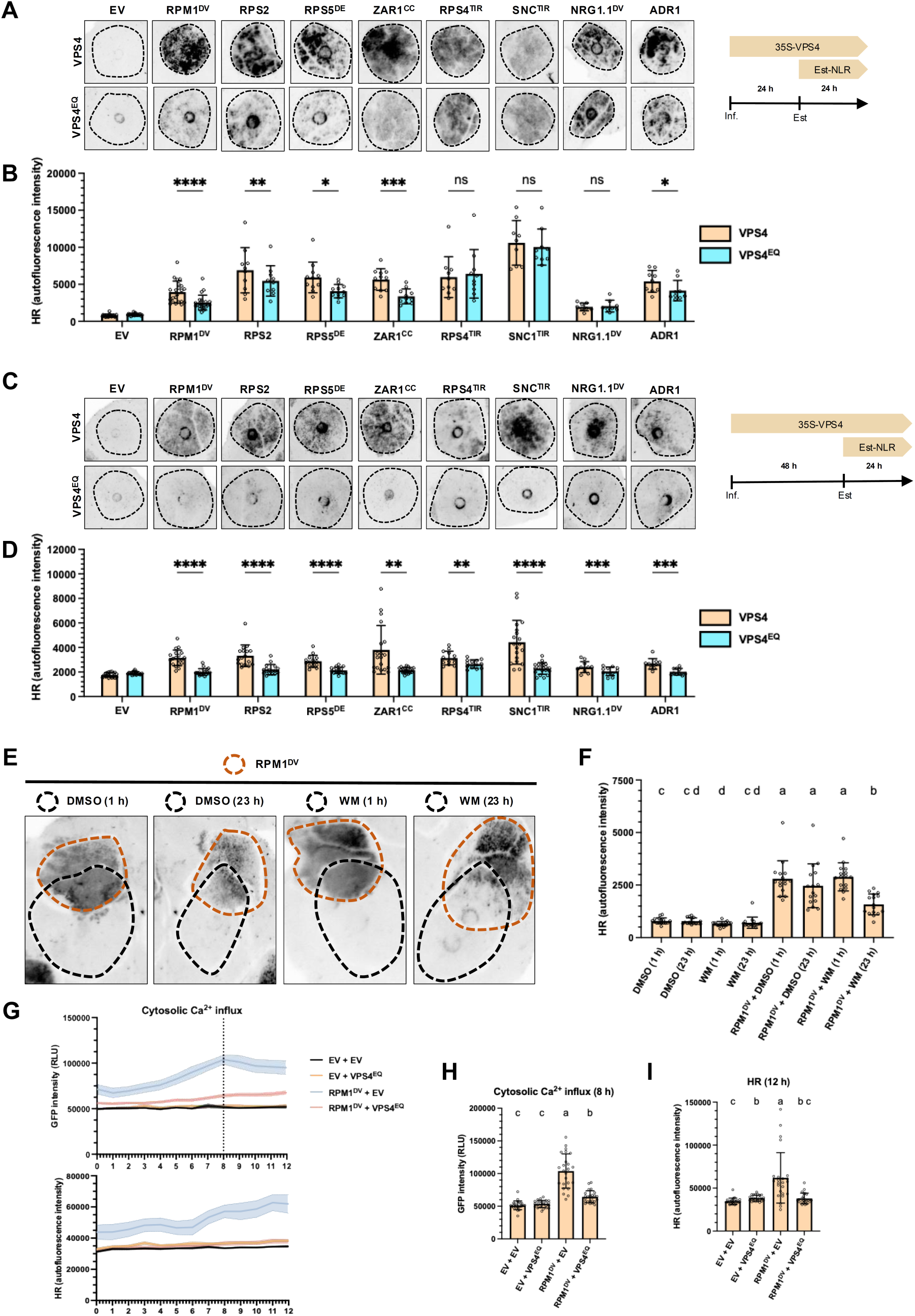
NLR-mediated HR is dependent on endomembrane trafficking. **(A and C)** Representative images of the inhibitory effects on HR by VPS4^EQ^ expression for 24 h **(A)** and 48 h **(C)** prior to estradiol induction of NLR expression in *N. benthamiana*. HR was visualized by red-light imaging 1 d after estradiol induction. **(B and D)** Quantification of HR by red-light imaging of **(A)** and **(C)**, respectively. Fluorescence intensity in the infiltrated spots was quantified in ImageJ. GFP (n_B_=22; n_D_=19), RPM1^DV^ (n_B_=22; n_D_=19), RPS2 (n_B_=10; n_D_=17), RPS5^DE^ (n_B_=11; n_D_=15), ZAR1^CC^ (n_B_=11; n_D_=19), RPS4^TIR^ (n_B_=10; n_D_=13), SNC1^TIR^ (n_B_=9; n_D_=19), NRG1.1^DV^ (n_B_=8; n_D_=11) and ADR1 (n_B_=9; n_D_=10). Significant differences were assessed by a two-tailed ratio paired Student’s t-test (ns, non-significant; *, *P* < 0.05; **, *P* < 0.01; ***, *P* < 0.001; ****, *P* < 0.0001). **(E)** Representative images of the effects of pre-treatments with Wortmannin (WM; 30 μM) or DMSO for 1 h or 23 h prior to estradiol induction of NLR expression. Red circles indicate where a construct expressing RPM1^DV^ was infiltrated, while black circles indicate where DMSO or WM were infiltrated. **(F)** Quantification of HR in **(E)** performed as in **(B)** and **(D)**. n=16. **(G)** Top, Ca^2+^-influx into the cytosol quantified in *N. benthamiana* GCaMP3 sensor line leaf discs in a microplate reader, where cytosolic Ca^2+^ was measured as GFP fluorescence intensities. Bottom, in-between HR quantification by red-light imaging in the same experiment. Constructs were *Agrobacterium* infiltrated, leaf discs were taken 24 h later and incubated in water overnight. The incubation water was then replaced by 50 μM estradiol, and subsequently GFP fluorescence intensities were recorded every hour. Thick lines represent average intensity, and the shading represents SE. n=24. (**H)** Quantification of cytosolic Ca^2+^ at 8 h after estradiol induction. **(I)** Quantification of HR at 12 h. n=24. For **(B)**, **(D)**, **(F)**, **(H)** and **(I)** bars represent means ± SD. Dots represent individual data points. For **(F)**, **(H)** and **(I)**, different letters represent statistical differences between treatments assessed by paired one-way ANOVA with Tukey’s honestly significant difference (HSD) (*P* < 0.05). These results are representative of the outcomes of at least three independent experiments.

Finally, we assessed Ca^2+^-influx into the cytoplasm, which is a hallmark for the initiation of ETI. As expected, we found the cytosolic Ca^2+^-level to be elevated upon RPM1^DV^-expression. Interestingly, this influx was significantly reduced when vacuolar traffic was blocked by expression of VPS4^EQ^ (Figure 4G and 4H). HR was quantified by red-light imaging in between the cytosolic Ca^2+^-readings in the same experiment (Figure 4G and 4I), whereby we confirmed that the microplate setup performed as the whole-leaf assays for scoring immune responses.

Taken together, our results suggest that blocking vacuolar traffic affects NLR-mediated HR in a quantitative and indirect manner. Blocking this pathway appears to impact HR activated by a wide range of NLR classes, although HR activated by CNLs and ADR1s is more sensitive than the one activated by TNLs and NRG1s. Additionally, the inhibition of the ETI immune signaling cascade seems to occur prior to the Ca^2+^-influx.

### The *Bh* fungus temporarily stalls the vacuolar pathway to suppress HR during infection

Having found that NLR-mediated HR is dependent on an active vacuolar pathway and that several *Bh* effectors inhibit this pathway and suppress MLA-activated HR, we wanted to test whether *Bh* itself does the same during the actual infection process. We again expressed the (SP)-RFP-AFVY vacuolar marker and the Golgi ST-YFP marker as tools in barley epidermal cells and inoculated with the fungus in a compatible interaction, using the susceptible cultivar Golden Promise. At 24 h after inoculation (hai), early in haustorium formation, both (SP)-RFP-AFVY and ST-YFP were observed in the ER, overlapping with the ERD2-mYFP and (SP)-mCherry-HDEL ER-markers, respectively. Importantly, this ER-stalling was only observed in attacked cells – independently of penetration success and formation of immature haustoria, but not in non-attacked neighboring cells of the same leaf. The ER-stalling is visible as reticular structures, not only near the attack sites, but also elsewhere in the cell and as a perinuclear ring (Figure 5A and Supplemental Figure 8A).

**Figure 5.**
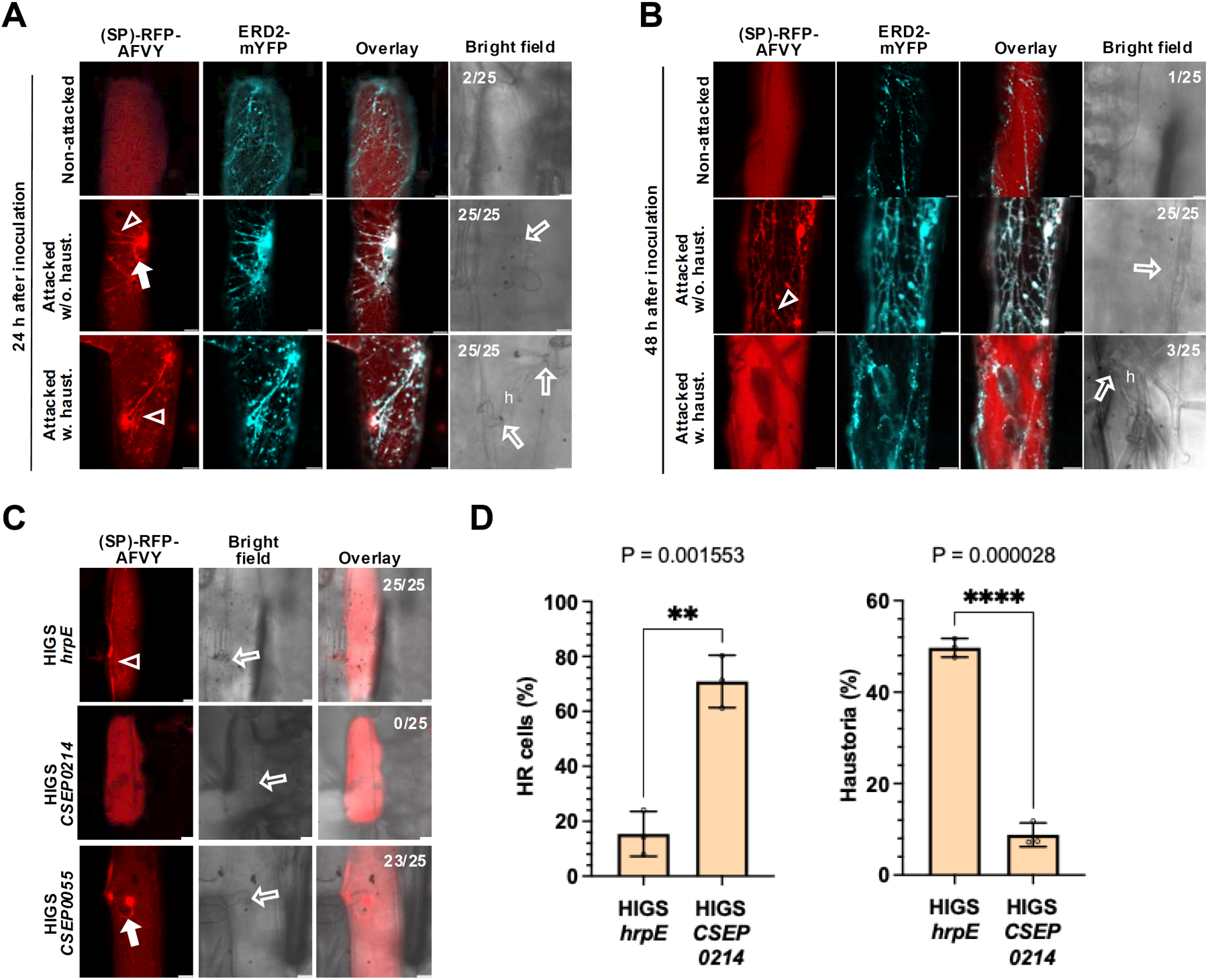
The *Bh* fungus temporarily stalls the endomembrane pathway to suppress HR during infection using effectors. **(A and B)** Vacuolar marker [(SP)-RFP-AFVY] stalled in ER with reticular signal overlapping with the ER marker, ERD2-mYFP, in barley leaf epidermal cells 24 **(A)** and 48 **(B)** hai with the virulent *Bh* isolate, C15. Numbers indicate proportions of cells showing ER-stalling as reticulate and perinuclear RFP signal in attacked cells without haustoria, successfully attacked cells with haustoria and non-attacked cells of the same leaf. **(C and D)** Host-induced gene silencing (HIGS) using hair-pin constructs against core effector CSEP0214, effector CSEP0055 not targeting endomembrane traffic (negative control), and the bacterial *hrpE* gene (negative control). **(C)** Attacked barley epidermal cells co-expressing (SP)-RFP-AFVY and HIGS constructs 24 hai with a virulent *Bh* isolate. Numbers indicate proportions of cells showing reticulate and perinuclear RFP signal. **(D)** In an experiment parallel to the one in **(C)**, the HR rates, visualized using propidium iodide, and the haustoria rates were scored in barley leaf epidermal cells co-expressing HIGS constructs and GUS as a marker for successful transformation, at 48 hai. Particle bombardment was carried out on 7-day-old susceptible barley cv. Golden Promise and inoculated 2 d later. Cells were imaged 24 **(A and C)** and 48 **(B and D)** hai. Bars represent means ± SD (n=3). Dots represent individual data points. Significant differences were assessed by a two-tailed unpaired Student’s t-test (**, *P* < 0.01; ****, *P* < 0.0001). Open arrowhead, reticular signal overlapping with ER marker. Closed arrow, perinuclear signal overlapping with the ER marker. Open arrows, *Bh* attack sites. h, haustoria. Scale bars, 10 μm. These results are representative of the outcomes of at least three independent experiments.

Surprisingly, when we studied ER-stalling in the same compatible interaction at 48 hai, we observed a pronounced difference. When the fungus successfully penetrated the epidermal cell and the haustorium had fully formed, the ER-stalling of (SP)-RFP-AFVY and ST-YFP was no longer observed at this later time-point. However, in attacked cells without haustoria, the ER-stalling remained as seen at 24 hai (Figure 5B and Supplemental Figure 8B). The dependence on haustorium formation for the release of the ER-stalling was confirmed in another susceptible barley line, Ingrid, and its *mlo-5*-resistant near-isogenic line (Freialdenhoven et al., 1996), in which immunity to *Bh* is manifested efficiently at the stage of penetration and formation of haustoria is prevented. In the *mlo-5*-resistant line, ER-stalling was observed in all attacked cells at both 24 and 48 hai. Whereas in the susceptible Ingrid, ER-stalling was only seen at 24 hai, while it was released at 48 hai after haustoria formation (Supplemental Figure 9), as observed previously.

To confirm that ER-stalling is caused by effectors targeting the vacuolar traffic to suppress HR when they are secreted by *Bh* during the attack, we took advantage of RNA interference (RNAi) of *BhCSEP0214* using host-induced gene silencing (HIGS) (Nowara et al., 2010). Silencing *BhCSEP0214* was enough to abolish the *Bh*-induced ER-stalling of the vacuolar marker (SP)-RFP-AFVY, which could instead be observed in the vacuole at 24 hai. In contrast, silencing the negative controls, *BhCSEP0055* and *hrpE*, had no effect on the ER-stalling of the vacuolar marker, which could still be observed at 24 hai (Figure 5C). In a parallel experiment, silencing *BhCSEP0214* caused a significant increase in the percentage of cells undergoing HR, from 15% to 71%, relative to the *hrpE* negative control. This corresponded to a decrease in the percentage of cells with haustoria (Figure 5D). In summary, these experiments indicate that during the infection process, the barley powdery mildew fungus uses the highly conserved core effector *Bh*CSEP0214 to target vacuolar traffic and prevent HR. It is possible that the remaining effectors serve a supporting role. Remarkably, we find the ER-stalling to be temporary and not occurring at a later time-point in successfully attacked cells, where haustoria have been fully formed.

To test whether the ER-stalling phenotype is restricted to barley leaf epidermal cells attacked by powdery mildew fungi, we examined wheat attacked by *B. graminis* f.sp. *tritici* (*Bgt*). In this compatible interaction, reticulate and perinuclear signals from the (SP)-RFP-AFVY marker, overlapping with the ERD2-mYFP ER marker, were also observed at 24 hai (Supplemental Figure 10A). At 48 hai, in attacked cells with a fully formed haustoria, the vacuolar marker was only observed at the vacuole (Supplemental Figure 10B). These observations parallel the results obtained in barley attacked by *Bh* and suggest that at least the essential powdery mildew fungi attacking grasses use the same strategy for suppressing immunity.

ER stress is commonly observed in plants during pathogen attack (Liu et al., 2025). Thus, we hypothesized the ER-stalling phenotype to correlate with ER stress during powdery mildew fungal attack. To analyze this, we investigated the transcript accumulation of genes commonly upregulated during the unfolded protein response (UPR) (Kim et al., 2018). We observed that the expression levels of many of these genes was already upregulated at 18 h after *Bh* inoculation in barley (Qian et al., 2023) (Supplemental Figure 11A and 11B) and at 12 h after *Bh* inoculation in partially immunocompromised *Arabidopsis* (Maekawa et al., 2012; Supplemental Figure 11C and 11D). This indicates that the ER-stalling induced by monocot-adapted powdery mildew fungi indeed correlates with occurrence of ER stress, and that the UPR is likely triggered when the fungus is suppressing vacuolar traffic in the host cell. Yet, this stress response apparently does not reach a sufficient level to trigger UPR-related programmed cell death (Pastor-Cantizano et al., 2024), which would otherwise be detrimental to the biotrophic powdery mildew fungus.

Dicot-adapted powdery mildew fungi also appear to benefit from blocking the host vacuolar pathway, as exemplified by *Golovinomyces orontii* infection in *Arabidopsis* (McRae et al., 2023; Nielsen et al., 2017; Schmidt et al., 2014). Schmidt et al. (2014) found the *G. orontii* effector *Go*BEC4 targets a barley ADP ribosylation factor-GTPase-activating protein, orthologous to the *Arabidopsis* AGD5. *Atagd5* mutants showed increased susceptibility to the powdery mildew fungus, *Erysiphe pisi*, and to the oomycete, *Hyaloperonospora arabidopsidis*. Additionally, the same study showed that Arabidopsis plants overexpressing *GoEC2* (homologous to *BhCSEP0214*) had increased susceptibility to *E. pisi*, but not to *G. orontii*, which was likely a reflection of the naturally high susceptibility already present in the *Arabidopsis*/*G*. *orontii* interaction (Schmidt et al., 2014). Strikingly, spray-induced gene silencing (SIGS) of *BhCSEP0214* homologues in *G. orontii* and *Erysiphe necator* significantly reduced fungal growth in *Arabidopsis* and *Vitis vinifera*, respectively (McRae et al., 2023).

### The *Bh* fungus secretes effectors to inhibit vacuolar traffic-suppressing effectors after haustorium formation

Our observation that the block of the vacuolar trafficking pathway is released only after haustoria have been fully formed suggested that this is a process controlled by the fungus and not by the plant host. *BhCSEP0214* is highly expressed at 6 hai and significantly downregulated closer to the time of haustorium formation (Supplemental Figure 12A; Sabelleck et al., 2025). However, *BhCSEP0162*, *BhCSEP0174*, and *BhCSEP0219* all seem to have higher expression closer to the time of haustorium formation (Supplemental Figure 12B-12D), which matches with the supposed timing of the release of the block of the vacuolar pathway. This contradictory result made us ask whether the fungus could secrete different effectors that interact with and inhibit vacuolar traffic-suppressing effectors after the haustorium is formed to effectively lift the block. To test this hypothesis, we used *Bh*CSEP0162 as a model and tested its interaction with other *Bh* effectors using Alphafold2-multimer (Evans et al., 2021; Jumper et al., 2021). Of the positive hits (Supplemental Data Table 2), *Bh*CSEP0142, *Bh*CSEP0198 and *Bh*CSEP0232 were confirmed to interact with *Bh*CSEP0162 by Y2H (Figure 6A). We then co-expressed these effectors with *Bh*CSEP0162 in *N. benthamiana* to test whether they can inhibit *Bh*CSEP0162-induced suppression of RPM1^DV^-mediated HR. This was indeed the case for *Bh*CSEP0142, that reverted the effect of *Bh*CSEP0162 on RPM1^DV^-mediated HR (Figures 6B and 6C). On the other hand, *Bh*CSEP0198 and *Bh*CSEP0232 could individually inhibit HR by themselves and therefore it was not possible to conclude whether they affected *Bh*CSEP0162-mediated HR inhibition (Figures 6D and 6G). Additionally, *Bh*CSEP0142 did not influence the HR-suppressing activity of *Bh*CSEP0214, indicating its specificity for *Bh*CSEP0162 (Figures 6H and 6I). Furthermore, expression of *Bh*CSEP0142, *Bh*CSEP0198, and *Bh*CSEP0232 individually did not induce ER-stalling of the vacuolar marker (SP)-RFP-AFVY in barley epidermal cells to the level of *Bh*CSEP0162 (Supplemental Figure 13). However, co-expression of *Bh*CSEP0142 led to lifting of the ER-block caused by *Bh*CSEP0162 (Supplemental Figure 13), likely explaining its effect on HR.

**Figure 6.**
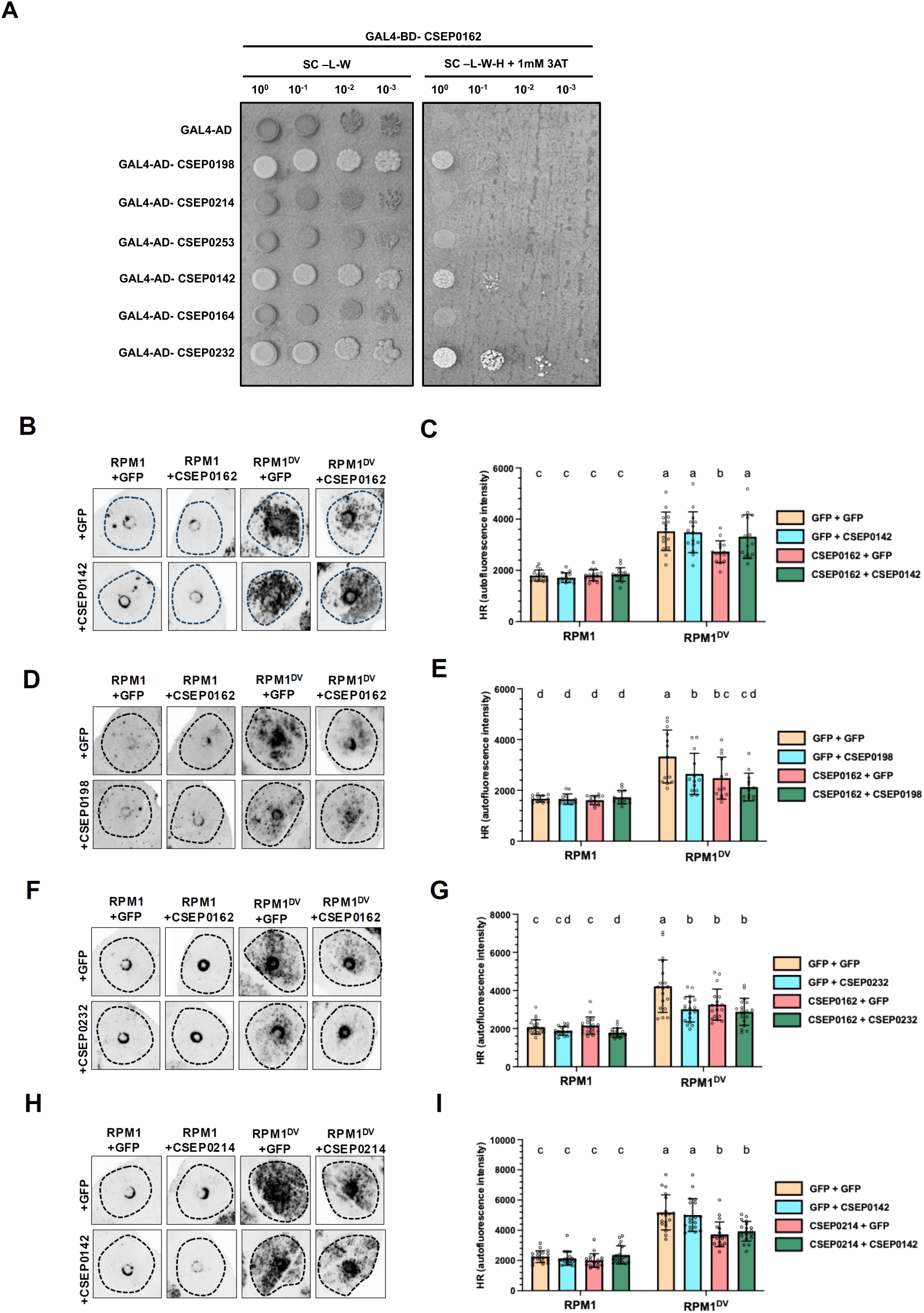
CSEP0142 prevents CSEP0162-mediated HR inhibition. **(A)** Binary Y2H interaction between CSEP0162 and putative interactor effectors identified through Alphafold2-multimer screening. **(B, D, F, and H)** Representative images of the effects on RPM1-mediated HR from the co-expression of CSEP0162 with CSEP0142 **(B)**, with CSEP0198 **(D)**, and with CSEP0232 **(F)**, and from the co-expression of CSEP0214 with CSEP0142 **(H)** after *Agrobacterium* infiltration in *N. benthamiana*. CSEPs were constitutively expressed for 48 h prior to induction of RPM1 using estradiol. HR was visualized by red-light imaging 1 d after estradiol induction. **(C, E, G, and I)** Quantification by red-light imaging of HR in **(B)**, **(D)**, **(F)**, and **(H)**. Average fluorescence intensity in the infiltrated areas was quantified using ImageJ. Error bars, ± SD; n=15 **(C)**; n=13 **(E)**; n=18 **(G)**; n=19 **(I)**. Dots represent individual data points. Different letters represent statistical differences between treatments assessed by paired three-way ANOVA with Tukey’s honestly significant difference (HSD) (P < 0.05). These results are representative of the outcomes of at least three independent experiments.

Overall, our results with *Bh*CSEP0162 suggest the possibility that *Bh* secretes effectors dedicated to inhibit vacuolar traffic-suppressing effectors to re-open the pathway after haustorium formation. Meanwhile, other effectors, such as *Bh*CSEP0198 and *Bh*CSEP0232, may inhibit HR through unknown mechanisms independently of blocking vacuolar traffic.

## DISCUSSION

Plant membrane trafficking is a prime target of many pathogens (Bhandari and Brandizzi, 2024; Nielsen, 2024; Yuen et al., 2023). Here we used ‘ER-stalling’ of a vacuolar marker as a proxy to study how powdery mildew fungi manipulate the vacuolar pathway. Surprisingly, the ER-stalling phenotype is conserved as far as the distantly related yeast fungus, suggesting the existence of a phenomenon that can be traced back to the ‘last eukaryotic common ancestor’ (LECA). An explanation for this phenomenon could be that the block of anterograde traffic to the vacuole/lysosome may disrupt retrograde trafficking pathways, thereby affecting earlier endomembrane traffic steps, eventually causing an ER UPR, which we for instance observe as the (SP)-RFP-AFVY ER-stalling. This is supported by the fact that knock-out of the retromer components, *VPS29* or *VPS35*, suppress vacuolar traffic and NLR-mediated resistance in Arabidopsis (Munch et al., 2015).

In the present study, we highlight the relevance of the vacuolar pathway in immunity by showing that the powdery mildew fungus uses effectors to stop this pathway to prevent ETI. It was recently observed that the vacuolar trafficking components, MON1, VPS18 and AMSH3, are required for activation of HR and immunity by the CNLs, MLA3, RPM1 and RPS2 (Liao et al., 2023; Sabelleck et al., 2025; Schultz-Larsen et al., 2018). In the present study, we further found that blocking vacuolar traffic has a general HR inhibitory effect, with HR activated by CNLs and ADR1s being more sensitive than TNL and NRG1-activated HR. This effect appears to be quantitative, since blocking vacuolar traffic for only a short time (1 h) prior to RPM1^DV^ activation was not sufficient to prevent HR. Similarly, blocking traffic for 24 h prior to TNL and NRG1.1 activation was also not sufficient to prevent HR. Together, these results indicate that vacuolar traffic is not required for NLR activation *per se*. Rather, it seems more likely that blocking vacuolar traffic progressively accentuates a cellular stress that is the actual cause of HR inhibition. However, we cannot exclude that vacuolar traffic may be required for elimination of a ‘negative regulator’ of HR prior to NLR activation. Different studies have previously shown that Arabidopsis mutants with defects in vacuolar traffic (*amsh3-4*, *vsr1 vsr6 vsr7* and *atg2*), either in the MVB or the autophagic pathways, also have hampered NLR-mediated HR, and this effect is stronger compared to the one observed in mutants with less severe traffic defects (*amsh3-3*, *vsr5 vsr6 vsr7*, *atg5*) (Munch et al., 2014; Schultz-Larsen et al., 2018; Zhu et al., 2025). This further emphasizes the quantitative nature of this phenomenon, and it is particularly striking that knocking out *NPR1* in the *atg2* and *atg5* backgrounds partially restores RPM1-induced HR (Munch *et al*., 2014). This suggests that HR does not directly require autophagy, but blocking autophagic traffic still suppresses HR, which seems to be a parallel to our results with MVB-mediated vacuolar traffic. We also observed that blocking vacuolar traffic affects the NLR-activated signaling events as early as the cytosolic Ca^2+^-influx stage, even though RPM1 protein accumulation and its subcellular localization were not affected. While Zhu et al. (2025) have recently shown that the vacuolar transport of hydrolytic enzymes and tonoplast-plasma membrane fusion are suppressed in *vsr* mutants during NLR-mediated ETI, it is important to consider that these processes likely happen downstream of cytosolic Ca^2+^-influx. How their data harmonizes with our finding that Wortmannin treatment for 1 h does not affect RPM1-triggered HR and that vacuolar traffic arrest impacts Ca^2+^-influx, awaits to be uncovered.

We provide evidence that powdery mildew fungi take advantage of the relationship between the vacuolar trafficking pathway and NLR-mediated resistance by actively impairing this pathway at early stages of infection. We found four unrelated *Bh* effectors that individually suppress *Mla*-locus-activated HR, likely by targeting vacuolar trafficking components when introduced into barley. It is important to note that *Mla*-specified resistance obviously works in an incompatible interaction despite expression of these vacuolar traffic-suppressing fungal effectors during *Bh* attack. This may reflect a quantitative balance of CNL-activation of HR and its inhibition by a block in vacuolar traffic. Thus, silencing the core effector, *Bh*CSEP0214, causes HR to be activated in an otherwise compatible interaction. This leads us to suggest that latent CNL-activation occurs in compatible interactions, and that a quantitative threshold determines whether NLRs can activate resistance after effector recognition. Removal of a core effector suppressing vacuolar traffic effectively enhances the immune response to make it reach this threshold, allowing HR to be activated in an otherwise compatible interaction. Meanwhile, in an incompatible interaction, MLA-activation plus the latent CNL-activation together trigger immunity sufficiently to surpass the threshold even when powdery mildew fungi secrete vacuolar traffic-suppressing effectors. Meanwhile, immunity can be suppressed by the overexpression of these same effectors, causing a stronger block of the vacuolar pathway. Overall, our results demonstrate the delicate balance between effective resistance and susceptibility to a pathogen (Figure 7A). It is also noteworthy that *Bh*CSEP0174 causes ER-stalling and suppresses HR, potentially by targeting AP1γ2, while it also is known as AVR_A9_, which leads to activation of HR when the CNL MLA9 is present (Saur et al., 2019). This may indicate that the timing of NLR activation relative to the block of vacuolar traffic may also play a role in the interaction, and that plants may pre-emptively counteract this pathogen strategy by early detection of effectors that target the vacuolar pathway.

**Figure 7.**
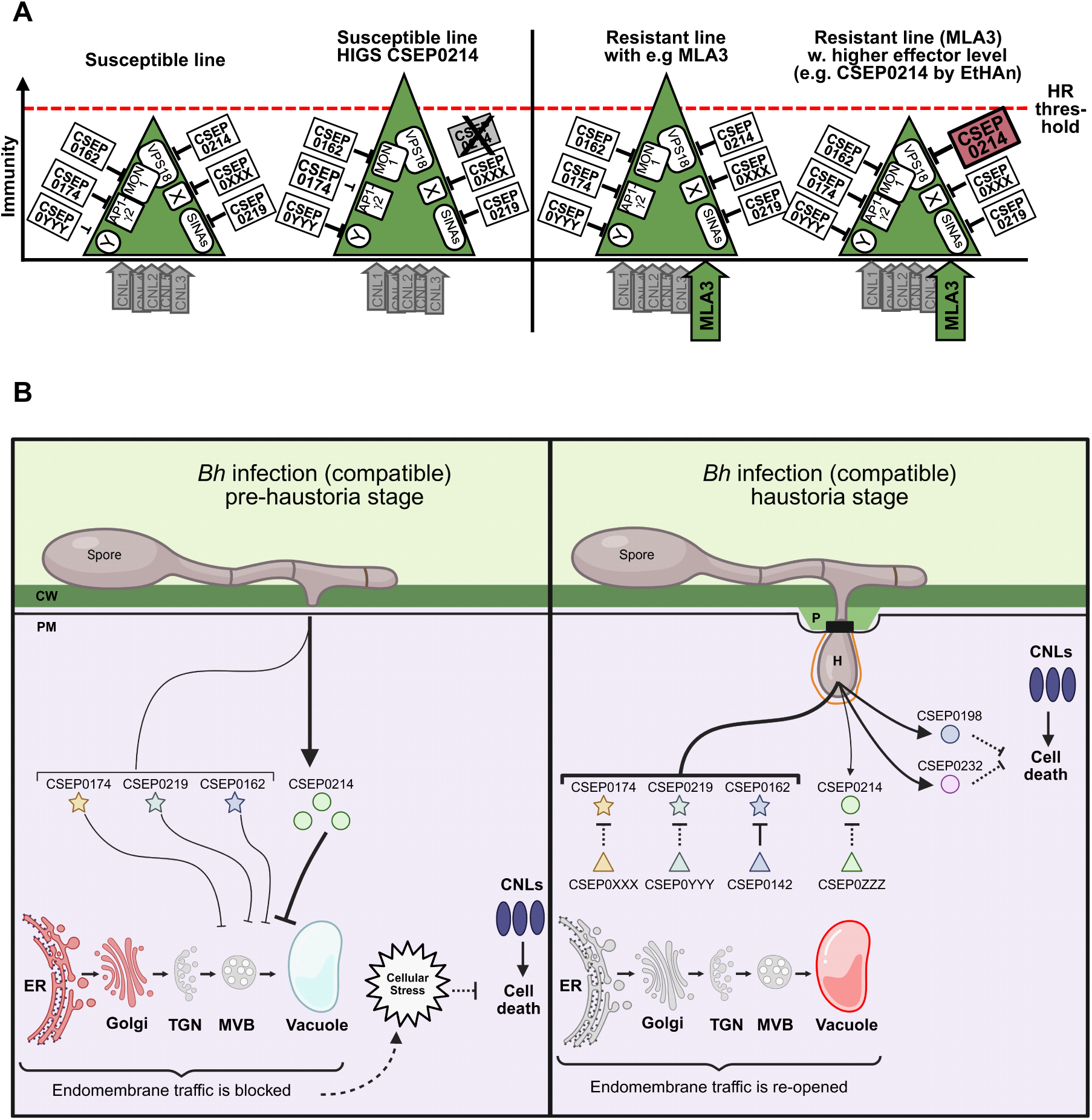
Models for the mechanism of vacuolar traffic suppression by *Bh* and how this affects NLR-mediated HR. **(A)** In a susceptible plant, powdery mildew fungi secrete several effectors that inhibit vacuolar traffic, suppressing latent CNL-activated immunity. If the core effector *CSEP0214* is silenced by HIGS, then the suppression is reduced, and the latent CNL-activated immunity reaches the threshold for HR manifestation. In a plant with e.g. MLA3-mediated resistance, the combined MLA3 and latent CNL-activated immunity is sufficient to reach the threshold for HR, even if vacuolar traffic is being suppressed. However, if the level of one of the vacuolar traffic-suppressing effectors in enhanced, for example by the EtHAn system, then the NLR suppression is increased, and the plants become susceptible even if MLA3 is activated. **(B)** During the actual infection, *Bh* inhibits vacuolar traffic at very early time-points, with CSEP0214 being the more prevalent effector in this process. When the haustoria are fully formed, the fungus re-opens the vacuolar trafficking pathway in the plant cell by simultaneously decreasing the expression of CSEP0214 and secreting a second set of effectors that inhibit the vacuolar traffic-suppressing effectors. The inhibition of CSEP0162 by CSEP0142 serves as an example of this. Additionally, *Bh* secretes more effectors such as CSEP0198 and CSEP0232 that inhibit NLR-mediated immunity independently of membrane traffic, thereby keeping NLR-mediated immunity suppressed even when the vacuolar pathway is re-opened (Created in BioRender. Deb, S. (2026) https://BioRender.com/avdu294).

Haustorium formation is critical for powdery mildew fungal growth. The fact that this pathogen targets and blocks vacuolar traffic at early stages of infection, and the fact that it can still grow when we artificially block the vacuolar pathway, suggests that formation of the extrahaustorial membrane (EHM), a process conducted by the plant, is traffic independent. This is consistent with our previous data, where haustoria formation takes place even when ER-to-Golgi traffic is inhibited (Kwaaitaal et al., 2017) and when the Rab7-GEF, MON1, is silenced (Liao et al., 2023). These observations indeed suggest that the EHM is formed independently of membrane trafficking. In the current study, we show that ER-stalling of (SP)-RFP-AFVY was released specifically in cells hosting haustoria at 48 hai, which suggested this to be a pathogen-controlled process. At this time-point the fungus starts using plant cell resources, and we postulate that only epidermal cells which have an active and well-functioning endomembrane trafficking system are good hosts for these obligate biotrophic pathogens. Although the results with *Bh*CSEP0162 are preliminary, they point to a model where powdery mildew fungi lift the block of the vacuolar trafficking pathway by reducing the expression of *Bh*CSEP0214 and by secreting a secondary set of effectors that inhibit the vacuolar traffic-suppressing effects of e.g. *Bh*CSEP0162. Additionally, effectors such as *Bh*CSEP0198 and *Bh*CSEP0232 can inhibit NLR-mediated HR independently of the vacuolar trafficking pathway, which can explain why the immune response does not restart after the block is lifted (Figure 7B). It could be that the fungus temporarily blocks vacuolar traffic as a mechanism to quickly suppress the general ETI response during the early time-points of infection, giving time for the pathogen to secrete additional effectors that inhibit HR independently of the vacuolar pathway. Why the fungus does not simply reduce the expression of *Bh*CSEP0162 and other vacuolar traffic-suppressing effectors as it does for *Bh*CSEP0214 by the time the haustorium is formed is still unknown. However, other groups have proposed that effector-effector interactions may serve to regulate and modulate effector function, leading to different outcomes in the host cell (Alcântara et al., 2019). It is possible that the interaction between *Bh*CSEP0162 with one of its effector interactors may provide it with a new function unrelated to the vacuolar pathway, and that this is why the fungus keeps its expression high even when vacuolar traffic should be unobstructed.

In the present study we benefitted from ER-stalling to directly visualize molecular and cellular processes essential for a plant-pathogen interaction. We believe this has revealed novel aspects of the complexity of these biological phenomena and further interlinked immunity and membrane trafficking. We have seen that ETI is surprisingly quantitative, that HR is only executed when a signaling threshold has been surpassed, and that effectors line up in suppressing immunity in a way that does not interfere with the pathogen taking control of the plant cell and persuade it to provide board and lodging.

## MATERIAL AND METHODS

### Plant and fungal material and growth conditions

Barley (*H. vulgare*), *N. benthamiana* and wheat (*Triticum aestivum*) plants were grown in a climate chamber under long-day conditions (16 h light/8 h dark), at 150 μE m^−2^ s^−1^ light intensity, 20°C (barley/wheat)/23°C (*N. benthamiana*), 60% relative humidity. Seven-day-old barley plants of the cultivar (cv.) Golden Promise, susceptible to the *Bh* isolate C15, were used for transient expression analysis, subcellular localization, and HIGS experiments. Near-isogenic lines P02, with the powdery mildew resistance gene *Mla3*, in the barley cv. Pallas background (Kølster et al., 1986) and the avirulent *Bh* isolate A6 (*avr_a1_, AVR_a3_*) were used to study effectors introduced by EtHAn. An *mlo-5*-carrying near-isogenic line of barley cv. Ingrid (Büschges et al., 1997; Freialdenhoven et al., 1996) and the *Bh* isolate C15 (*AVR_a1_, avr_a3_*) provided an interaction with strong penetration resistance. *N. benthamiana* plants were used for protein expression analysis, subcellular localization and cell death assays. Wheat cv. Sharki and a local uncharacterized virulent powdery mildew fungal (*B. graminis* f.sp. *tritici*, *Bgt*) isolate was used for subcellular localization.

### Yeast strains and growth conditions

Yeast (*Saccharomyces cerevisiae*) strain BY4741 (Euroscarf acc. no. Y00000) and the *vps18* mutant in the BY4741 background (Euroscarf acc. no. Y04105; genotype: BY4741; MATa; *his3*Δ1; *leu2*Δ0; *met15*Δ0; *ura3*Δ0; YLR148w::*kanMX4*) were used for subcellular localization experiments. Yeast cells were transformed following the standard LiAc protocol (Gietz and Woods, 2002) with (SP)-RFP-AFVY or (SP)-RFP-QRPL in Gateway® cloning vectors for yeast (Alberti et al., 2007), along with pTPI1-GFP-HDEL (Addgene id: ZJOM144).

### Yeast two-hybrid next-generation interaction screens

Yeast two-hybrid next-generation interaction screens (Y2H-NGIS) were performed as previously described (Elmore et al., 2023; Li et al., 2024; Velásquez-Zapata et al., 2023). Three independent replicates of CSEP0174, CSEP0219 and non-select controls were used as baits for three independent replicates of batch Y2H-NGIS, using a three-frame cDNA prey library of 1.1 x 10^7^ primary clones generated from our 6-point infection time course (Velásquez-Zapata et al., 2022). Libraries representing each replicate were sequenced to 20 million reads by Genewiz from Azenta (Morrisville, NC, USA). Candidate interactors were quantitatively ranked via Y2H-SCORES criteria, comprising enrichment, specificity, and in-frame ensemble (Supplemental Data Table 1; Velásquez-Zapata et al., 2021). Interacting prey fragments were determined using alignments obtained via NGPINT (Smith et al., 2025; Velásquez-Zapata et al., 2023), brought in-frame, and cloned into the prey vector (pDEST-AD) for binary Y2H confirmation tests (Dreze et al., 2010). These were performed by plating three dilutions of each bait/prey mating on media selecting for diploids (SC-LW) or interactions (SC-LWH + 0.5 mM 3-AT). Yeast colonies were photographed 4 days after plating.

### Cloning

The coding sequences for *Bh* effectors were synthesised by ThermoFisher Scientific, NLR coding sequences were amplified from Arabidopsis gDNA using primers listed in Supplemental Data Table 3 and cloned via Gateway BP reactions into pDONR201 or pDONR207, or by TOPO cloning reaction into pENTR/D-TOPO. The coding sequences were subsequently transferred to destinations vectors (see Supplemental Data Table 4) using Gateway LR reactions according to the manufacturer’s protocol (Supplemental Data Table 5). pMDC7-mYFP-GWY was made by digesting pMDC7 (Curtis and Grossniklaus, 2003) with XhoI and inserting mYFP, cloned from pMDC7-GWY-mYFP-HA (Gao *et al*., 2011), by Gibson Assembly (New England Biolabs GmbH, Frankfurt, Germany).

### Transient expression and RNA interference in barley

Transient expression of untagged effector proteins was carried out using the pUbi::GWY vector (Shen et al., 2007). Transient-induced gene silencing of barley genes and host-induced gene silencing (HIGS) of powdery mildew fungal genes (Nowara et al., 2010) were performed by transiently transforming cells with over-expression and hair-pin constructs. DNA constructs were transformed into epidermal cells of the abaxial side of leaves from 7-day-old barley seedlings by particle bombardment, as previously described (Douchkov et al., 2005). Bombardment of 1-µm gold particles coated with the DNA constructs (see Supplemental Data Table 5) was performed using the PDS-1000/He particle delivery system (Bio-Rad, Basel, Switzerland), mounted with a hepta-adapter. Visualization of transformed cell was either performed via detection of fluorescent proteins by confocal imaging or by light microscopy of β-GUS-stained cells (Douchkov et al., 2005).

### Agroinfiltration

*Agrobacterium tumefaciens* strain GV3101 (pMP90) transformed with binary vector constructs (Karimi et al., 2002; Nakagawa et al., 2007) (see Supplemental Data Table 5) were grown in LB media with selected antibiotics, and infiltrated into leaves of 3-4-week-old *N. benthamiana* plants with a 1-mL needleless syringe after resuspension in infiltration buffer (10 mM MgCl_2_, 10 mM MES pH 5.6, 200 µM acetosyringone), mixed as indicated in Supplemental Data Table 6. For estradiol induction, the plant expression vectors pMDC7, pMDC7-GWY-mYFP-HA (Gao et al., 2011) or pMDC7-mYFP-GWY were used, and plants were sprayed with 20 μM estradiol solution at the indicated times. Samples for protein expression analysis were collected 5 h after estradiol induction. Wortmannin (Cayman Chemical, Michigan, USA), 30 μM in DMSO, was infiltrated the same way at the indicated times.

### Bimolecular fluorescence complementation (BiFC) assay

For BiFC assays, leaves of 4 week-old *N. benthamiana* plants were infiltrated with *A. tumefaciens* strains carrying the genes of interest tagged with either the N- or C-terminal part of red fluorescence protein (RFP). After 2 days, leaf discs for all tested interactions were visualised by confocal laser scanning microscopy.

### Confocal imaging

The adaxial side of barley and *N. benthamiana* leaves, and yeast cultures, were visualized using a Stellaris 8 (Leica, Wetzlar, Germany) confocal laser scanning microscope. Plant and yeast cells were visualized using 20x and 40x oil objectives, respectively. Laser excitation/detection wavelengths for mYFP/GFP and mCherry/RFP were 508/513-575 nm and 587/596-644 nm, respectively.

### EtHAn-mediated introduction of effectors for studying of impact on immune responses

The *Pseudomonas fluorescence* Effector-to-Host Analyzer (EtHAn) strain expressing a type III secretion system (Thomas et al., 2009) was transformed with selected constructs in the pEDV6-GWY vector (Fabro et al., 2011) (see Supplemental Data Table 5), and was infiltrated into barley P02 (*Mla3*) leaves with a 1-mL needleless syringe, followed by inoculation with the avirulent *Bh* isolate, A6. Cell death was visualized by Trypan Blue staining followed by destaining with chloral hydrate (Koch and Slusarenko, 1990). Coomassie blue staining was used to visualize fungal structures.

### HR quantification

HR cell death responses in *N. benthamiana* were quantified at the indicated time-points using red light imaging (Villanueva et al., 2021). Briefly, infiltrated leaves were visualized in a ChemiDoc MP imaging system (Bio-Rad, Basel, Switzerland), model Universal Hood III, equipped with the Green LED Module kit no. 1708284. Samples were excited in the ‘green epi illumination’ setting, and the 605/50 emission filter was used. Average fluorescence intensity was quantified using Fiji (Schindelin et al., 2012). Single cell HR responses of barley epidermal cells were either assayed following Trypan Blue staining (Koch and Slusarenko, 1990), or following propidium iodide staining at a concentration of 10 µg/ml (Truernit and Haseloff, 2008).

### Cytosolic Ca^2+^-influx assay

The *N. benthamiana* GCaMP3 Ca^2+^ sensor line was used to measure the levels of cytosolic Ca^2+,^ and the assay was performed as previously described, with slight modifications (DeFalco et al., 2017). All the different construct combinations were agroinfiltrated in different spots on the same leaf to reduce experimental variability. T7-RPM1^DV^ was expressed under an estradiol inducible promoter using the pMDC7 expression vector. Leaf discs measuring 0.5 cm in diameter were taken 24 h after agroinfiltration and equilibrated in 200 μL ddH_2_O overnight, in a Greiner 96-well flat black plate. Then, the incubation solution was replaced by 200 μL ddH_2_O containing 50 μM estradiol and measurements were performed every hour starting from that time-point on a Clariostar Plus plate reader (BMG LABTECH, Ortenberg, Germany). GCaMP3 was excited at 475-505 nm and its fluorescence emission was detected at 510-560 nm. Parallel to this, HR was quantified by exciting the samples at 494-570 nm and detecting fluorescence emission at 605-649 nm.

### Protein purification and western blot analysis

Samples for protein expression were homogenized in protein extraction buffer [50 mM Tris-HCl, pH 8.0, 50 mM NaCl, 1 mM EDTA, 0.5% NP40, 1 mM dithiothreitol, Complete Mini protease inhibitor cocktail (Roche, Mannheim, Germany)]. Protein concentration was determined by the Bradford reaction (Bradford, 1976). SDS-PAGE and immunoblot analysis were done using standard procedures. For antibodies, see Supplemental Data Table 7.

### Transcriptomic analysis

A list of genes commonly upregulated during UPR in Arabidopsis has been previously published (Kim et al., 2018). A BLAST search was carried out with their protein sequences against the barley proteome assembly (MorexV3) using EnsemblPlants (Yates et al., 2022), with an E-value<0.05. When multiple homologues were identified, the one with the closest homology was chosen as a representative. Publicly available transcriptomic datasets (accession numbers PRJNA835302 and GSE39463) (Maekawa et al., 2012; Qian et al., 2023), were utilized for the analysis of ER stress during powdery mildew infection. The dataset PRJNA835302 was analyzed by performing independent filtering based on gene variance using the "genefilter" package (version 1.91.0) (Gentleman et al., 2025), and differential expression analysis was performed using the "limma" package (Ritchie et al., 2015). For the GSE39463 dataset, low count genes were filtered and differential expression analysis was performed using the "DEseq2" package (version 1.49.2) (Love et al., 2014). Further details regarding the datasets, gene identification, and statistical analysis can be found in Supplemental Data Table 8.

### Statistical analysis and reproducibility

For experiments involving quantification, plotting and statistical analysis was performed on GraphPad Prism 10. Information about the type of statistical test performed, description of the plot elements, and number of repetitions are mentioned in the figure legend for the respective experiments. Data used for plotting and statistical analysis, as well as all the statistical analysis parameters chosen, are provided in the Source Data file. Uncropped blots for Supplementary Figure 6A are also shown in a Source Data file.

## Data availability

All relevant data are within the manuscript and its Supplemental Data Files. Raw Y2H-NGIS sequence reads are deposited at the NCBI Gene Expression Omnibus (GEO) under the accession numbers GSE299565 (CSEP0174) and GSE299566 (CSEP0219). Full instructions for the NGPINT alignment software are located in the GitHub repository https://github.com/Wiselab2/NGPINT_V3 and Zenodo https://doi.org/10.5281/zenodo.15256036. Container images are hosted on Sylabs (https://cloud.sylabs.io/library/schuyler/ngpint/ngpint) and Dockerhub (https://hub.docker.com/r/schuylerds/ngpint) for integration into high-throughput and cloud-computing workflows. R code, ReadMe and user instruction files for the Y2H-SCORES software are provided at GitHub (https://github.com/Wiselab2/Y2H-SCORES/tree/master/Software).

## ACKNOWLEDGMENTS

We thank Dr. Stefan Kusch for providing the counts table for the PRJNA835302 RNAseq dataset. The research was supported by the Novo Nordisk Fonden grant NNF19OC0056457 (PlantsGoImmune) to HTC, Villum Fonden grants 00028131 and 00050260 to HTC, Horizon Europe Marie Skłodowska-Curie Actions fellowship 101104193 to SD, Fulbright - Minciencias 2015 & Schlumberger Faculty for the Future fellowships to VVZ, USDA-ARS Postdoctoral Research Associateship and USDA-NIFA-ELI Postdoctoral Fellowship 2017-67012-26086 to JME, and National Science Foundation - Plant Genome Research Program grant 13-39348, USDA-National Institute of Food and Agriculture grant 2020-67013-311, Oak Ridge Institute for Science and Education (ORISE) under U.S. Department of Energy agreement 60-5030-2-004, and USDA-Agricultural Research Service projects 3625-21000-067-00D and 5030-21220-068-000-D to RPW. The funders had no role in study design, data collection and analysis, decision to publish, or preparation of the manuscript. Mention of trade names or commercial products in this publication is solely for the purpose of providing specific information and does not imply recommendation or endorsement by the USDA, NIFA, ARS, or the National Science Foundation. USDA is an equal opportunity provider, employer and lender. Figure 7B was created in BioRender.com.

## AUTHOR CONTRIBUTIONS

SD performed all cell biological (co-)localization studies in yeast and *in-planta*. SD and BS performed yeast mutant analyses. JPAP performed the calcium influx assay and localization study of RPM1 *in-planta*. JPAP, SD, BR and AC performed HR assays. VVZ, JME, and GF performed the Y2H-NGIS prey library screens, informatic analysis, and Y2H binary confirmation. XL performed yeast interaction studies. JPAP performed the transcriptomic data statistical analysis. PD performed the first VPS4^EQ^/(SP)-RFP-AFVY experiment. HTC conceived the study and supervised the project. HTC, SD and JPAP drafted and edited the manuscript; RPW directed the Y2H-NGIS projects and edited the manuscript.

## SUPPLEMENTAL FIGURE LEGENDS

**Supplemental Figure 1. Vacuolar marker is stalled in the ER when the endomembrane pathway is inhibited in yeast.**

**Supplemental Figure 2. Binary yeast two-hybrid and bimolecular fluorescence complementation (BiFC) confirmation of interactions identified by Y2H-NGIS.**

**Supplemental Figure 3. Barley powdery mildew effectors targeting the endomembrane trafficking pathway cause ER-stalling of vacuolar and Golgi markers in barley leaf epidermal cells.**

**Supplemental Figure 4: Bh effectors interfere with endomembrane traffic to the vacuole in Nicotiana benthamiana.**

**Supplemental Figure 5. RNA interference of barley endomembrane trafficking targets of powdery mildew effectors cause ER-stalling of vacuolar marker.**

**Supplemental Figure 6. Protein level and localization of RPM1^DV^ appear unchanged upon expression of VPS4^EQ^.**

**Supplemental Figure 7. Multiple negative regulators of vacuolar traffic inhibit NLR-mediated HR.**

**Supplemental Figure 8. The *Bh* fungus temporarily stalls the ST-YFP Golgi marker in the ER in attacked cells with haustoria.**

**Supplemental Figure 9. The *Bh* fungus releases the ER-stalling of the vacuolar marker at 48 h after inoculation only in barley cells with haustoria.**

**Supplemental Figure 10. The wheat powdery mildew fungus (*Bgt*) causes a vacuolar marker to be temporarily stalled in the ER in attacked cells of wheat by 24 hai and releases the ER-stalling at 48 hai in cells with haustoria.**

**Supplemental Figure 11. The expression levels of genes commonly upregulated during ER stress peaks at the onset of haustoria formation in barley and partially immunocompromised Arabidopsis attacked by the *Bh* fungus.**

**Supplemental Figure 12. Expression patterns of vacuolar trafficking-suppressing CSEPs during barley infection by *Bh*.**

**Supplemental Figure 13. Release of barley powdery mildew effector CSEP0162-mediated ER-stalling by its interacting effector, CSEP0142.**

**Supplemental Data Table 1. Yeast two-hybrid next-generation interaction screen for barley targets of *Bh*CSEP0219 and *Bh*CSEP0174.**

**Supplemental Data Table 2. AF2-multimer modelling of CSEP0162 dimerization**

**Supplemental Data Table 3. Primers used in this study.**

**Supplemental Data Table 4. DNA vectors used in this study.**

**Supplemental Data Table 5. DNA constructs used in this study.**

**Supplemental Data Table 6. Information regarding plant infiltration experiments.**

**Supplemental Data Table 7. Antibodies used in this study.**

**Supplemental Data Table 8. ER stress transcriptomic analysis.**

**Source Data for Figures 3, 4, 5, and 6 and for Supplemental Figures.**

**Source Data for Supplemental Figure 6.**

