## Supplemental Figure for "Powdery mildew fungi block plant vacuolar traffic to suppress immunity"

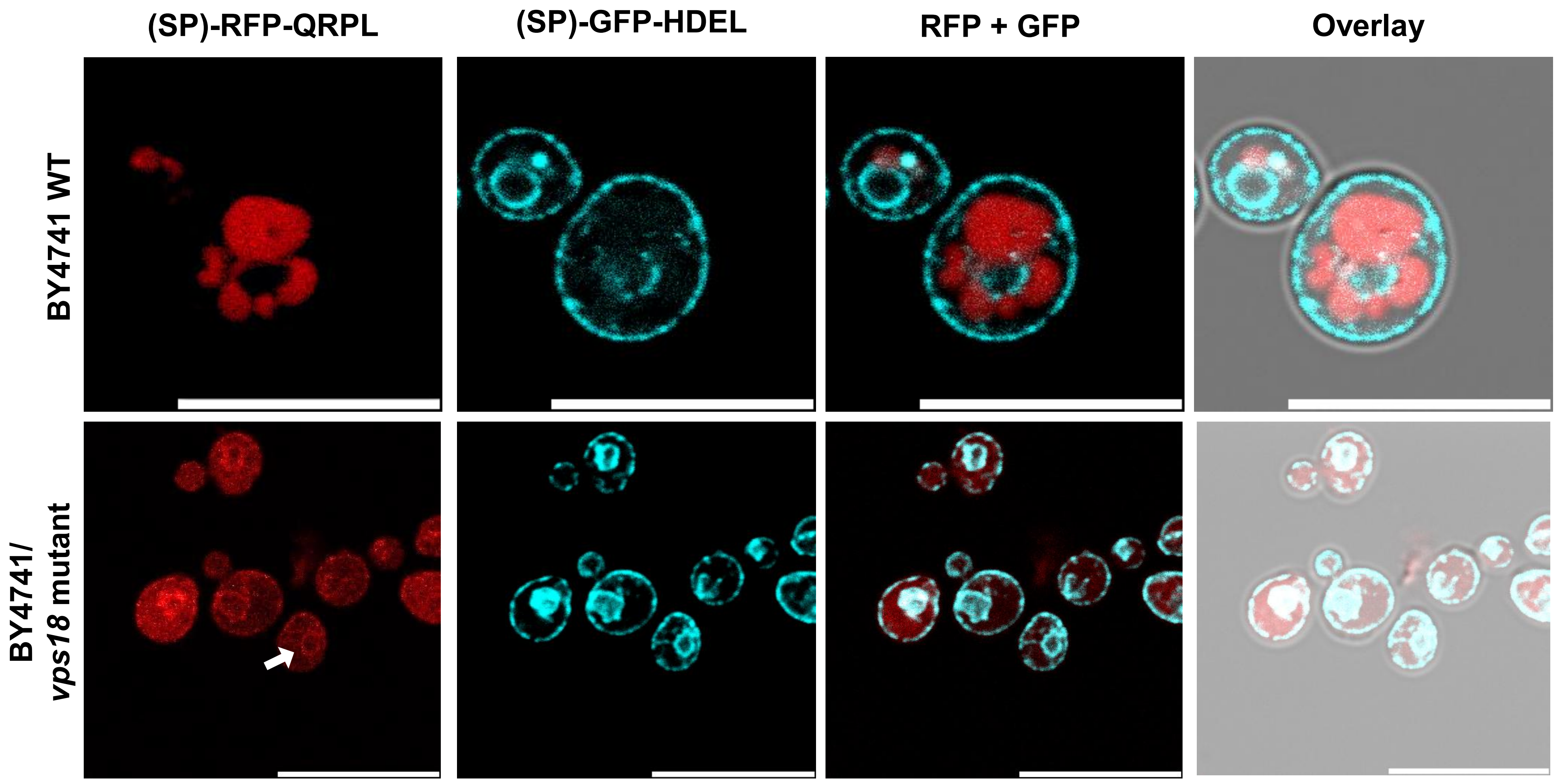


**Supplemental Figure 1. Vacuolar marker is stalled in the ER when the endomembrane pathway is inhibited in yeast.** Vacuolar marker [(SP)-RFP-QRPL] is stalled in the ER, with reticular and perinuclear signal overlapping with the ER marker [(SP)-GFP-HDEL] in stably transformed *vps18* mutant, but not in WT, of yeast strain BY4741. Closed arrow, perinuclear signal overlapping with the ER marker. Open arrowhead/box, reticular signal overlapping with ER marker. Scale bars, 10 µm. These results are representative of the outcomes of at least three independent experiments.

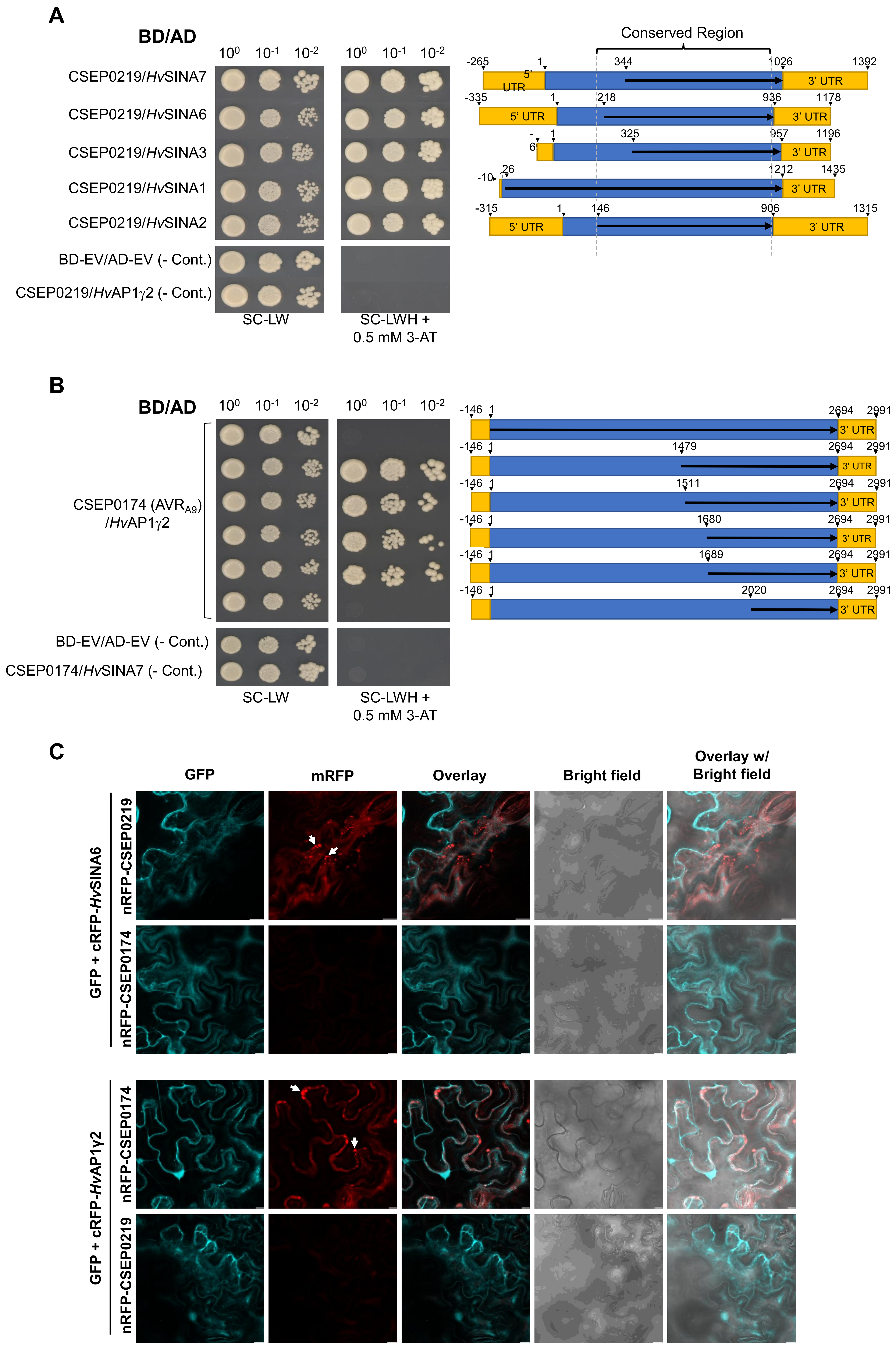
A
BD/AD
CSEP0219/HvSINA7
CSEP0219/HvSINA6
CSEP0219/HvSINA3
CSEP0219/HvSINA1
CSEP0219/HvSINA2
BD-EV/AD-EV (- Cont.)
CSEP0219/HvAP1γ2 (- Cont.)
SC-LW
SC-LWH + 0.5 mM 3-AT
Conserved Region
5' UTR
3' UTR
B
CSEP0174 (AVRA9) /HvAP1γ2
CSEP0174/HvSINA7 (- Cont.)
C
GFP
mRFP
Overlay
Bright field
Overlay w/ Bright field
GFP + cRFP-HvSINA6
nRFP-CSEP0219
nRFP-CSEP0174
GFP + cRFP-HvAP1γ2

**Supplemental Figure 2. Binary yeast two-hybrid and bimolecular fluorescence complementation (BiFC) confirmation of interactions identified by Y2H-NGIS. (A and B)** Interactions of five *Hv*SINA E3 ligases, identified as preys using CSEP0219 as bait, and of the adaptor protein, *Hv*AP1γ2, identified as prey using CSEP0174 as bait (see Supplemental Data Table 1), were tested by binary Y2H according to Dreze et al. (2010). CSEP0219 and CSEP0174 were fused to the C-terminus of the Gal4 transcription factor binding domain, while the indicated coding sequences (arrows) were used to fuse the preys to the C-terminus of the Gal4 transcription factor activation domain. **(A)** The five SINA E3 ligases interacting with CSEP0219 are numbered according to the barley chromosomal location of their encoding genes. **(B)** The C-terminal, but not the full-length, *Hv*AP1γ2 showed interaction with CSEP0174. CSEP0174/HvSINA7, CSEP0219/*Hv*AP1γ2 and BD/AD empty vectors were included as negative/specificity controls. **(C)** Interaction of CSEP0219 and *Hv*SINA6, and of CSEP0174 and *Hv*AP1γ2 shown by bimolecular fluorescence complementation in leaf epidermal cells of *Nicotiana benthamiana*. Reverse combinations used as negative/specificity controls. Arrowhead, endosome-like structures. Scale bars, 10 μm.

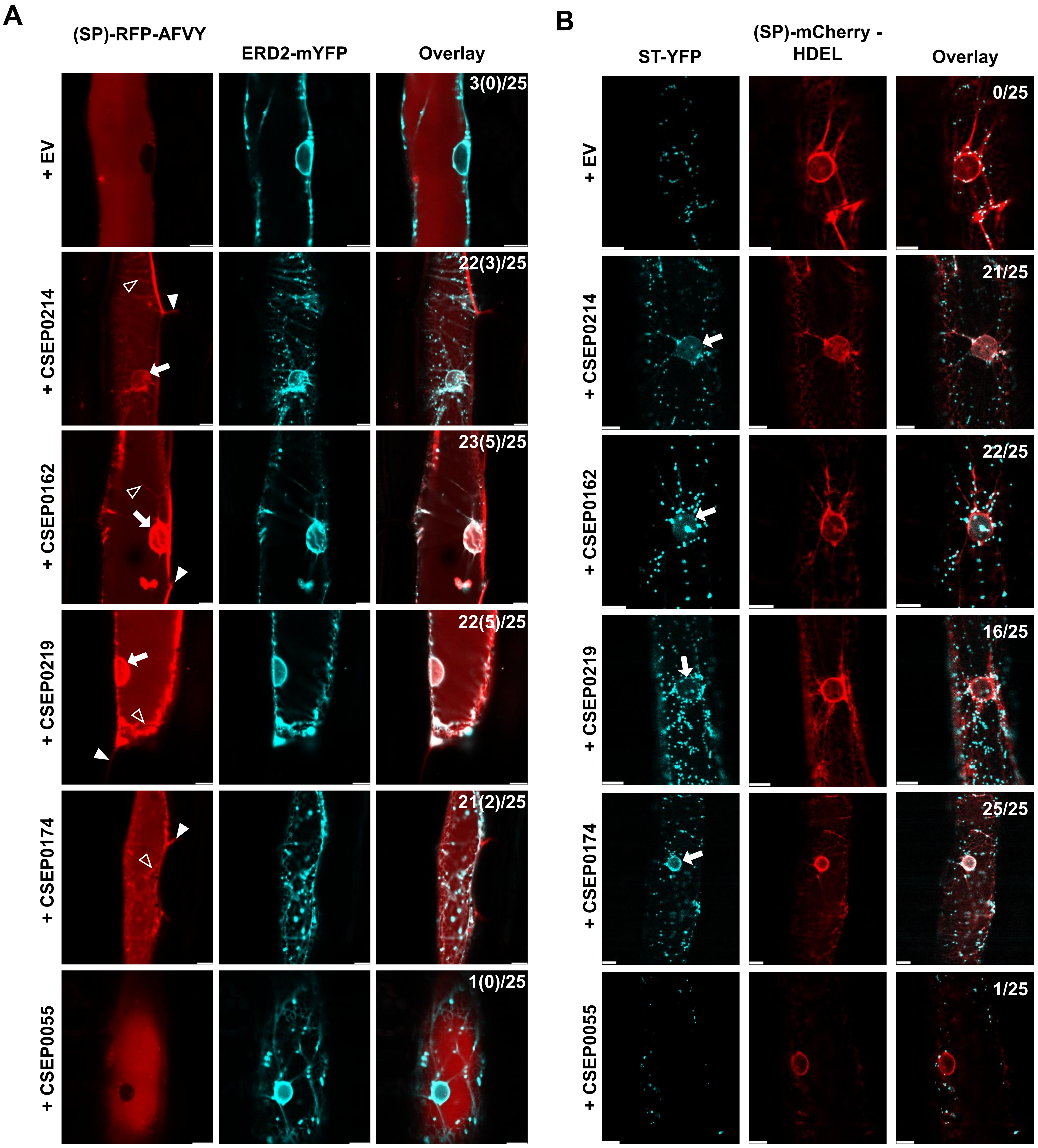


**Supplemental Figure 3. Barley powdery mildew effectors targeting the endomembrane trafficking pathway cause ER-stalling of vacuolar and Golgi markers in barley leaf epidermal cells. (A)** Vacuolar marker, (SP)-RFP-AFVY, stalled in ER with reticular and perinuclear signal overlapping with the ER markers, ERD2-mYFP. Numbers indicate proportions of epidermal cells showing ER-stalling and secretion (in brackets) of the vacuolar marker (closed arrowhead). **(B)** Golgi marker, ST-YFP, stalled in ER with perinuclear signal overlapping with the ER markers, (SP)-mCherry-HDEL. CSEP0055, negative control effector not targeting endomembrane traffic. EV, empty vector. Numbers indicate proportions of epidermal cells showing ER-stalling. Particle bombardment was carried out on 7-day-old barley *cv.* Golden Promise plants and cells were imaged 2 days later. Open arrowhead, reticular signal overlapping with ER marker. Closed arrow, perinuclear signal overlapping with the ER marker. Closed arrowhead, extracellular signal observed between two cells neighboring the transformed cell. Scale bars, 10 µm. These results are representative of the outcomes of at least three independent experiments.

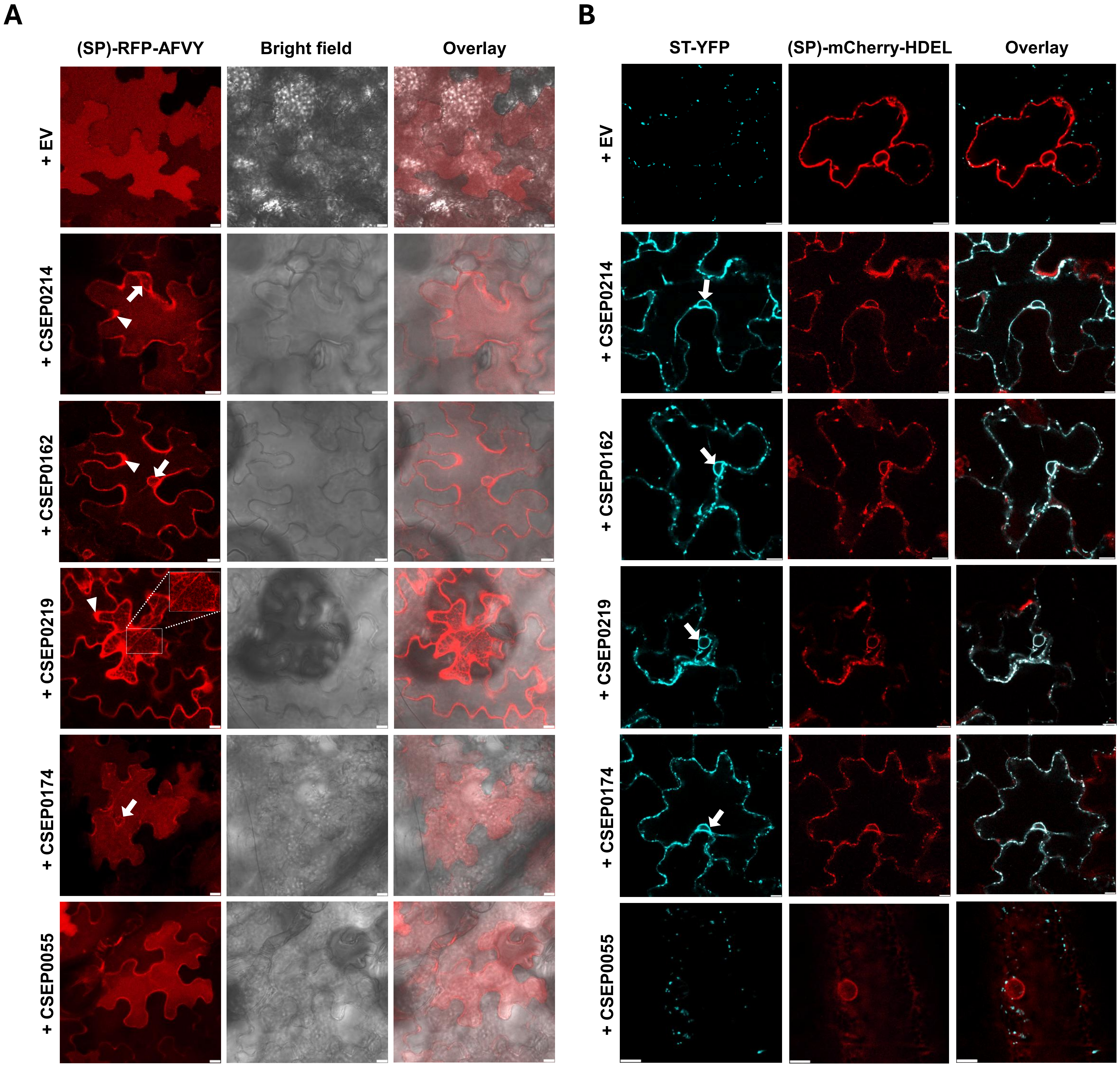


**Supplemental Figure 4: *Bh* effectors interfere with endomembrane traffic to the vacuole in *Nicotiana benthamiana*. (A)** Vacuolar marker, (SP)-RFP-AFVY, show reticular, perinuclear and/or extracellular signal in *N. benthamiana* leaf epidermal cells when co-expressed with *Bh* effectors. **(B)** Golgi marker, ST-YFP, co-localize with the ER marker, (SP)-mCherry-HDEL, at the perinuclear ring when co-expressed with *Bh* effectors. *N. benthamiana* transformed by *Agrobacterium* infiltration. Empty vector (EV) and CSEP0055 were used as negative control. Box, reticular signal. Closed arrow, perinuclear signal overlapping with the ER marker. Closed arrowhead, extracellular signal. Scale bar, 10 μm. These results are representative of the outcomes of at least three independent experiments.

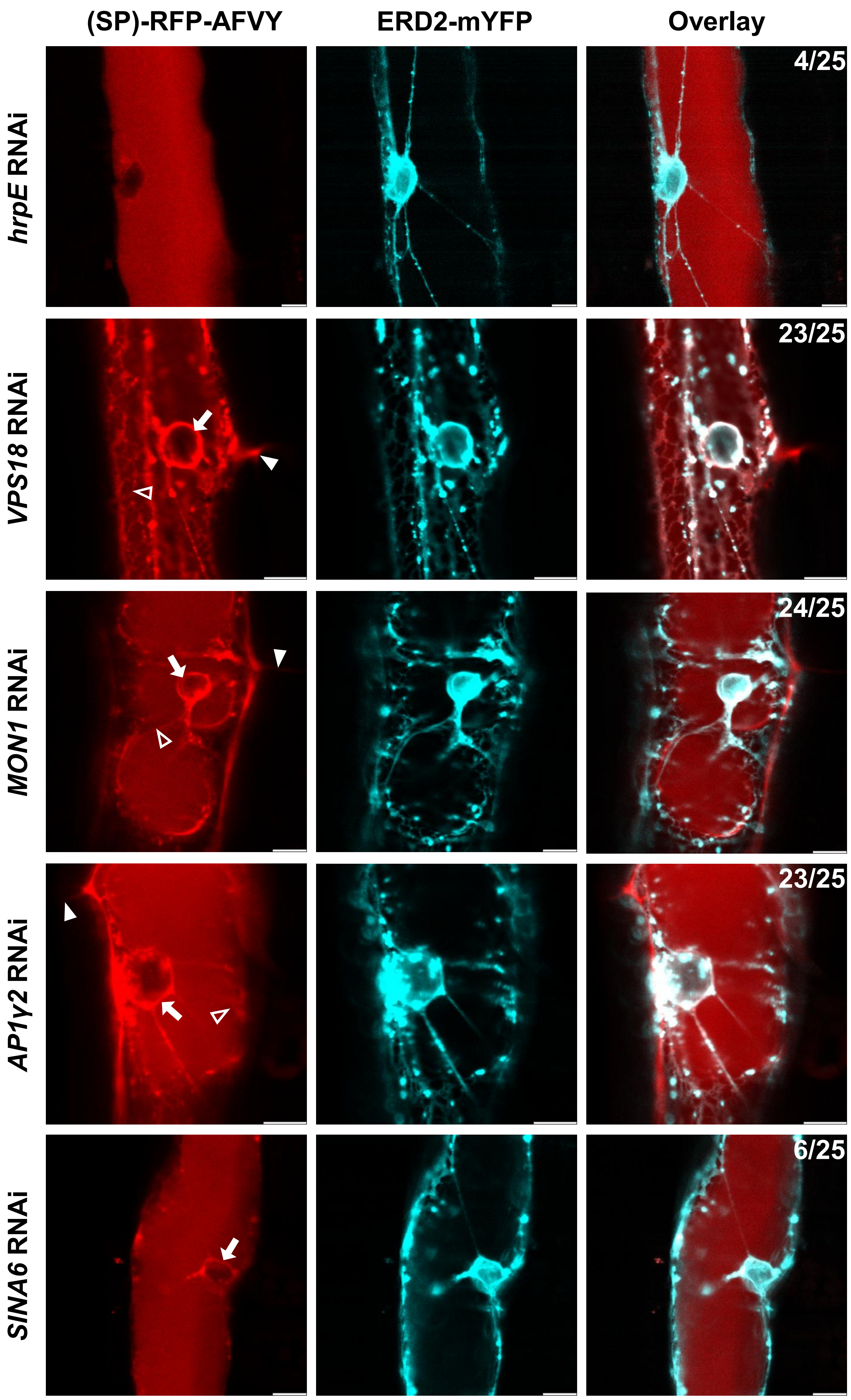


**Supplemental Figure 5. RNA interference of barley endomembrane trafficking targets of powdery mildew effectors cause ER-stalling of vacuolar marker.** RNAi hair-pin constructs of powdery mildew effector targets caused (SP)-RFP-AFVY stalling in ER with perinuclear and reticular signals overlapping with the ER marker, ERD2-mYFP. Hair-pin construct of bacterial *hrpE* used as negative control. Numbers indicate proportions of leaf epidermal cells showing ER stalling. Particle bombardment was carried out on 7-day-old barley *cv.* Golden Promise plants and cells were imaged 2 days later. Closed arrow, perinuclear signal overlapping with the ER marker. Open arrowhead, reticular signal overlapping with ER marker. Closed arrowhead, extracellular signal observed between two cells neighboring the transformed cell. Scale bars, 10 µm. These results are representative of the outcomes of at least three independent experiments.

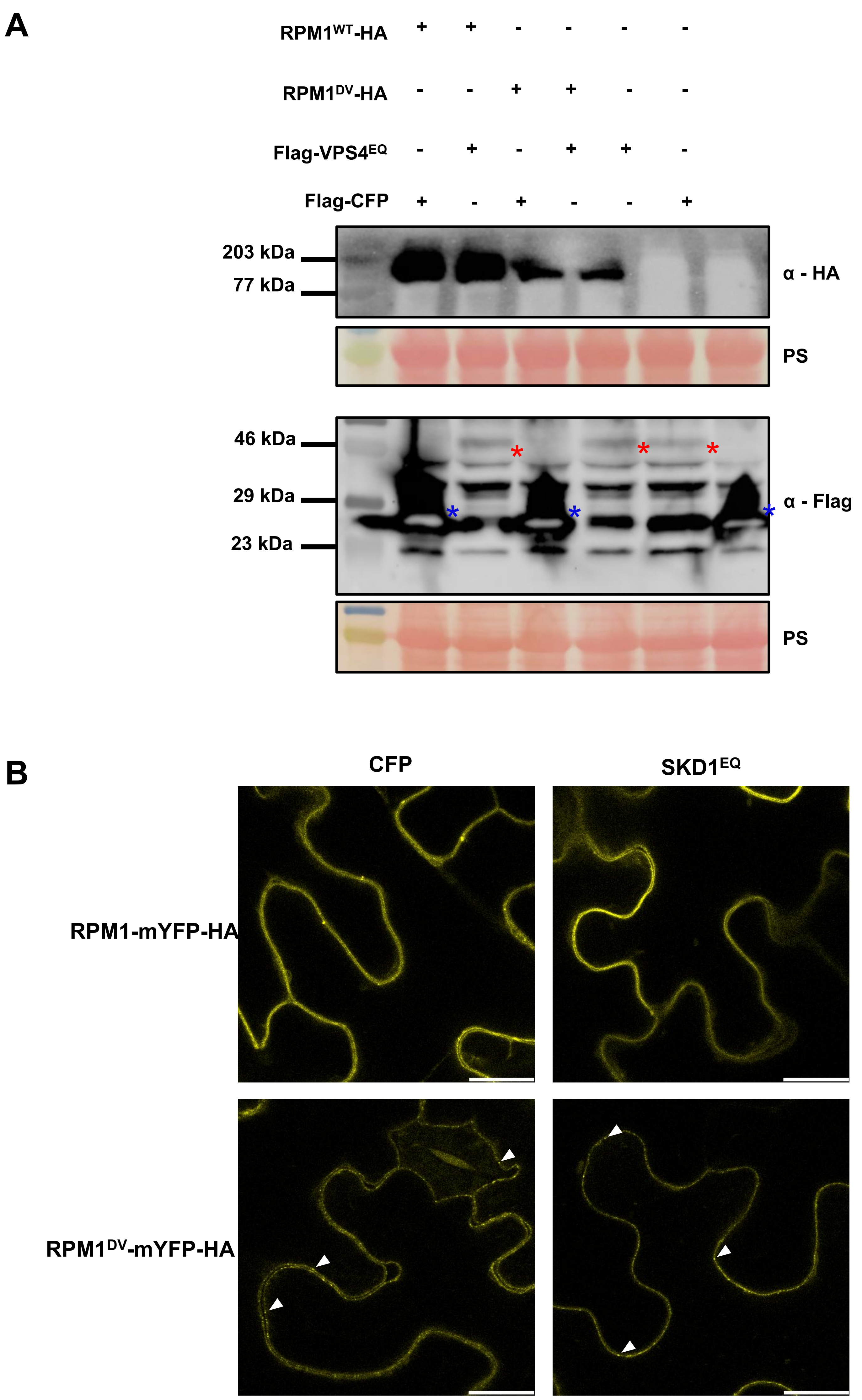


**Supplemental Figure 6. Protein level and localization of RPM1$^{DV}$ appear unchanged upon expression of VPS4$^{EQ}$.** Fusion proteins RPM1-mYFP-HA, RPM1$^{DV}$-mYFP-HA, Flag-VPS4$^{EQ}$ or Flag-CFP were co-expressed in different combinations after *A. tumefaciens*-mediated transient gene expression in leaves of *N. benthamiana* plants. **(A)** The leaves were sprayed with estradiol 48 hours post infiltration, and samples were collected for protein extraction 5 hours later. Total protein extract was used for SDS-PAGE and immunoblot analysis using α-HA and α-Flag antibodies. On the left, molecular masses of marker proteins are indicated. Ponceau staining (PS) was used to demonstrate equal amount of total protein on the blots. Red stars indicate bands of Flag-VPS4$^{EQ}$ and blue stars indicate bands of Flag-CFP. **(B)** The leaves were sprayed with estradiol 48 hours post infiltration and RPM1 subcellular localization was observed 5 hours later by confocal laser scanning microscopy on the mYFP channel. Closed arrows, aggregated RPM1$^{DV}$ at the plasma membrane. Scale bars, 20 µm. These results are representative of the outcomes of two independent experiments.

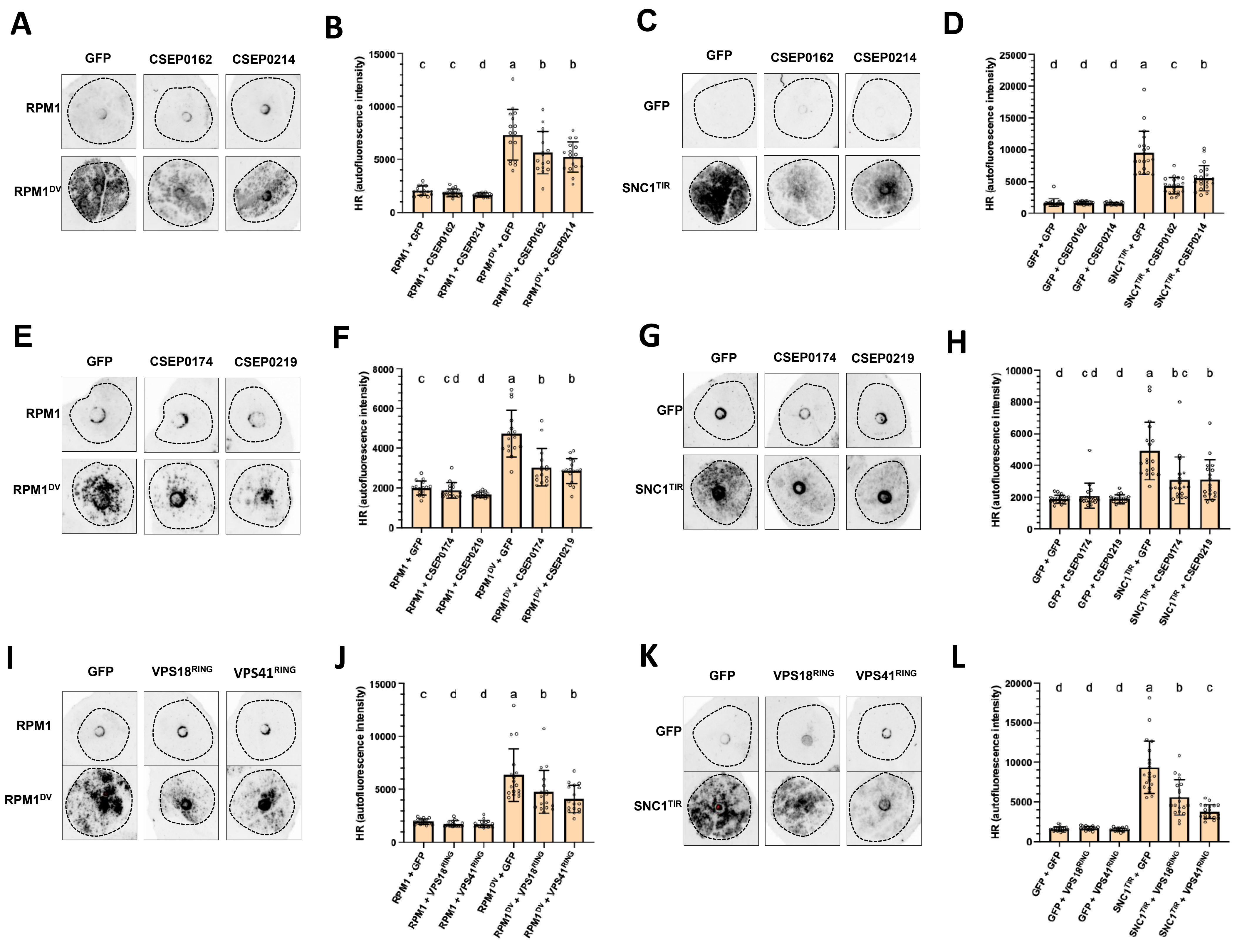


**Supplemental Figure 7. Multiple negative regulators of vacuolar traffic inhibit NLR-mediated HR. (A, C, E, G, I and K)** Representative images of HR inhibition mediated by *Bh*CSEP0162 and *Bh*CSEP0214 **(A and C)**, *Bh*CSEP0174 and *Bh*CSEP0219 **(E and G)**, and VPS18$^{RING}$ and VPS41$^{RING}$ **(I and K)**. HR was induced either by the CNL RPM1$^{DV}$ **(A, E, and I)** or the TNL SNC1$^{TIR}$ **(C, G, and K)**. The experiments were performed in *N. benthamiana* and HR was visualized by red-light imaging 3 d after *A. tumefaciens*-mediated transient gene expression and 1 d after estradiol-mediated induction of NLR expression. Black circles denote the area of infiltration. **(B, D, F, H, J, and L)** Quantification of HR by red-light imaging in **(A)**, **(C)**, **(E)**, **(G)**, **(I)** and **(K)**. Fluorescence intensity in the infiltrated areas was quantified using ImageJ. Bars represent means ±SD; n=16 **(B)**; n=20 **(D)**; n=16 **(F);** n=18 **(H)**; n=17 **(J)**; n=20 **(L)**. Dots represent individual data points. Letters represent statistical differences between treatments assessed by paired one-way ANOVA with Tukey's honestly significant difference (HSD) ($P < 0.05$). These results are representative of the outcomes of at least two independent experiments.

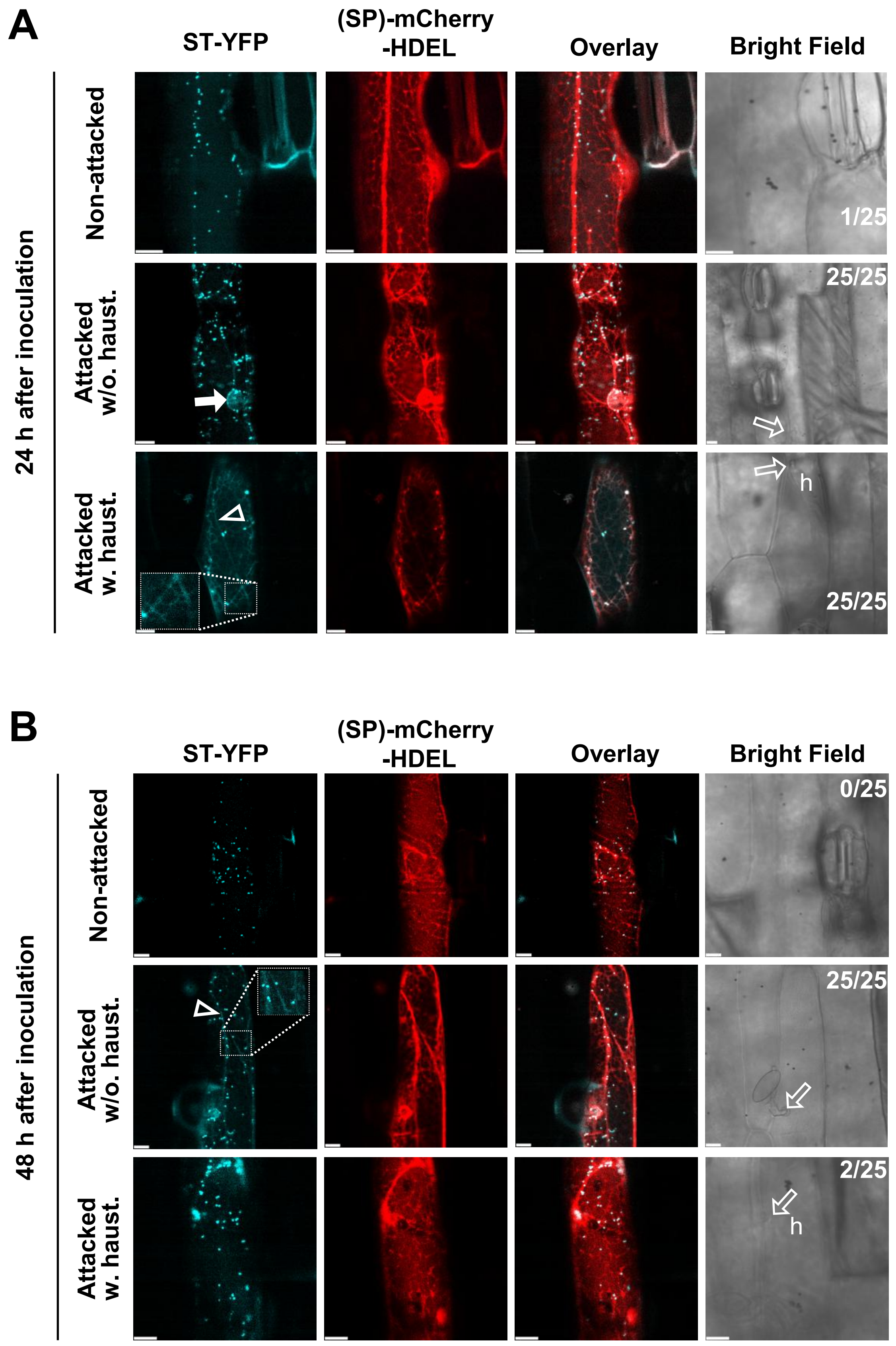


**Supplemental Figure 8. The *Bh* fungus temporarily stalls the ST-YFP Golgi marker in the ER in attacked cells with haustoria. (A and B)** ST-YFP Golgi marker stalled in ER with reticular (insert) and perinuclear signal overlapping with the ER markers, (SP)-mCherry-HDEL, in barley leaf epidermal cells 24 **(A)** and 48 **(B)** hai with a virulent *Bh* isolate. Numbers indicate proportions of cells showing ER stalling in non-attacked cells, non-successfully attacked cells without haustoria and successfully attacked cells with haustoria. Open arrowhead/box, reticular signal overlapping with ER marker. Closed arrow, perinuclear signal overlapping with the ER marker. Open arrows, *Bh* attack sites. h, haustoria. Scale bars, 10 µm. These results are representative of the outcomes of at least three independent experiments.

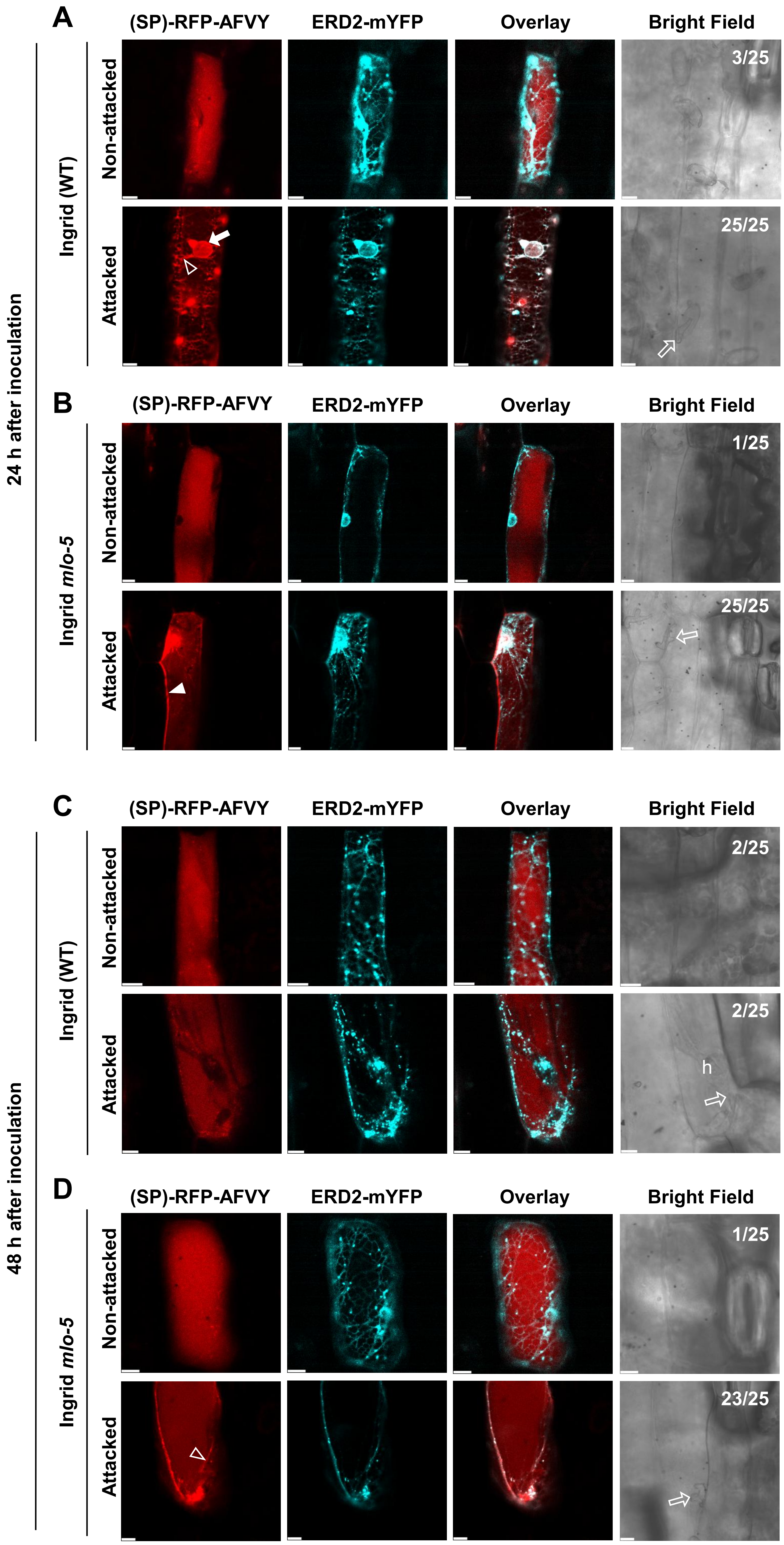
A
B
C
D
(SP)-RFP-AFVY
ERD2-mYFP
Overlay
Bright Field
24 h after inoculation
48 h after inoculation
Ingrid (WT)
Ingrid mlo-5
Non-attacked
Attacked
3/25
25/25
1/25
25/25
2/25
2/25
h
1/25
23/25

**Supplemental Figure 9. The *Bh* fungus releases the ER-stalling of the vacuolar marker at 48 h after inoculation only in barley cells with haustoria. (A-D)** Vacuolar marker, (SP)-RFP-AFVY, stalled in ER with reticular signal overlapping with the ER marker, ERD2-mYFP, in barley leaf epidermal cells of wildtype Ingrid attacked by *Bh* without penetration at 24 hai **(A)** and resistant cv. Ingrid *mlo-5* attacked by *Bh* without penetration at 24 hai **(B)** and at 48 hai **(D)**. However, in susceptible Ingrid (WT) cells, where haustoria had formed, ER-stalling was released at 48 hai **(C)**. Numbers indicate proportions of cells showing ER-stalling. Open arrowhead, reticular signal overlapping with ER marker. Closed arrow, perinuclear signal overlapping with the ER marker. Closed arrowhead, extracellular signal. Open arrows, *Bh* attack sites. h, haustoria. Scale bars, 10 μm. These results are representative of the outcomes of at least three independent experiments.

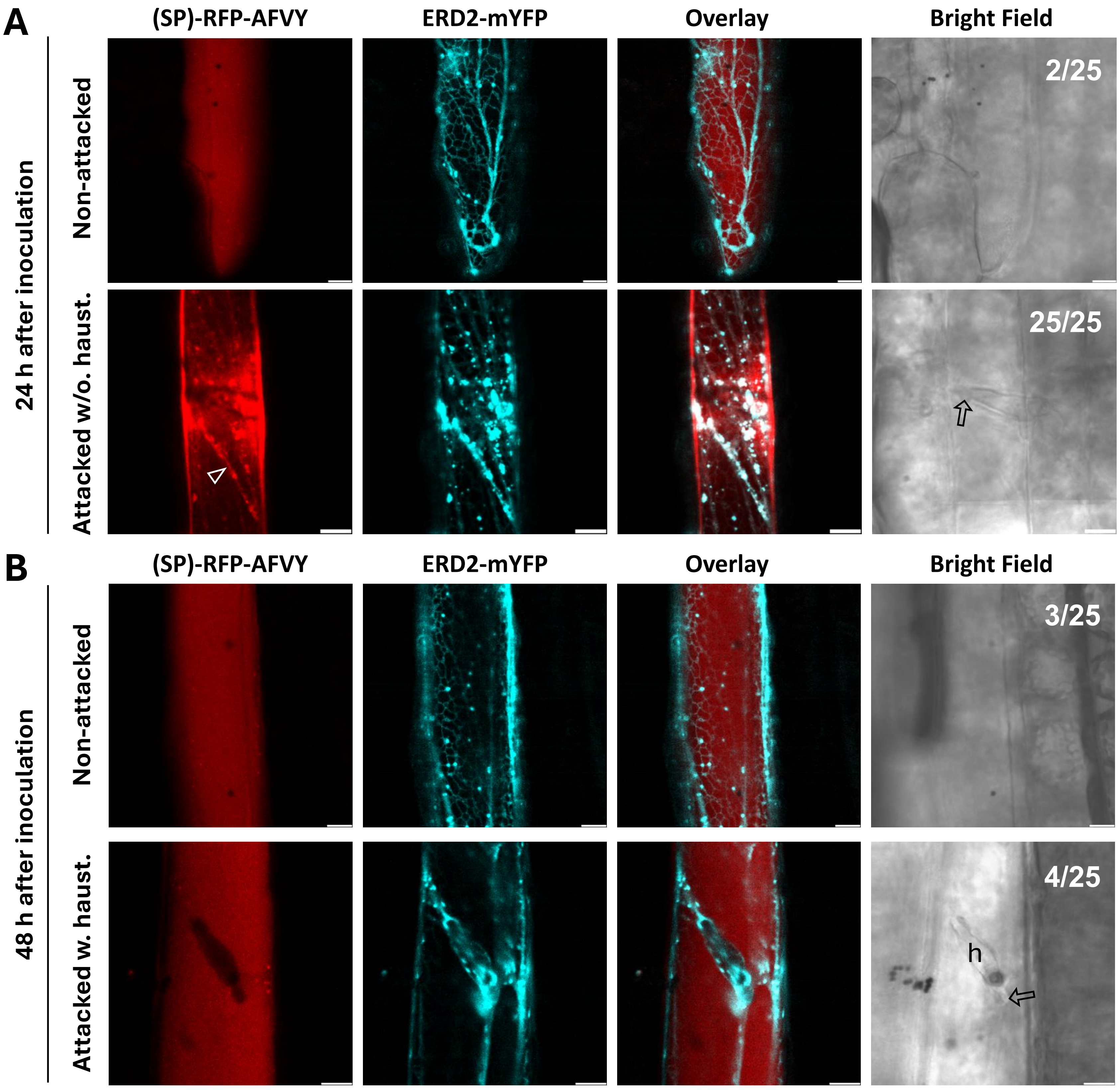


**Supplemental Figure 10. The wheat powdery mildew fungus (*Bgt*) causes a vacuolar marker to be temporarily stalled in the ER in attacked cells of wheat by 24 hai and releases the ER-stalling at 48 hai in cells with haustoria.** Vacuolar marker, (SP)-RFP-AFVY, stalled in ER with reticular signal overlapping with the ER markers, ERD2-mYFP, in wheat cv. Sharki leaf epidermal cells. Numbers indicate proportions of cells showing ER-stalling. Open arrowhead, reticular signal overlapping with ER marker. Open arrows, *Bgt* attack sites. h, haustoria. Scale bars, 10 µm. These results are representative of the outcomes of at least three independent experiments.

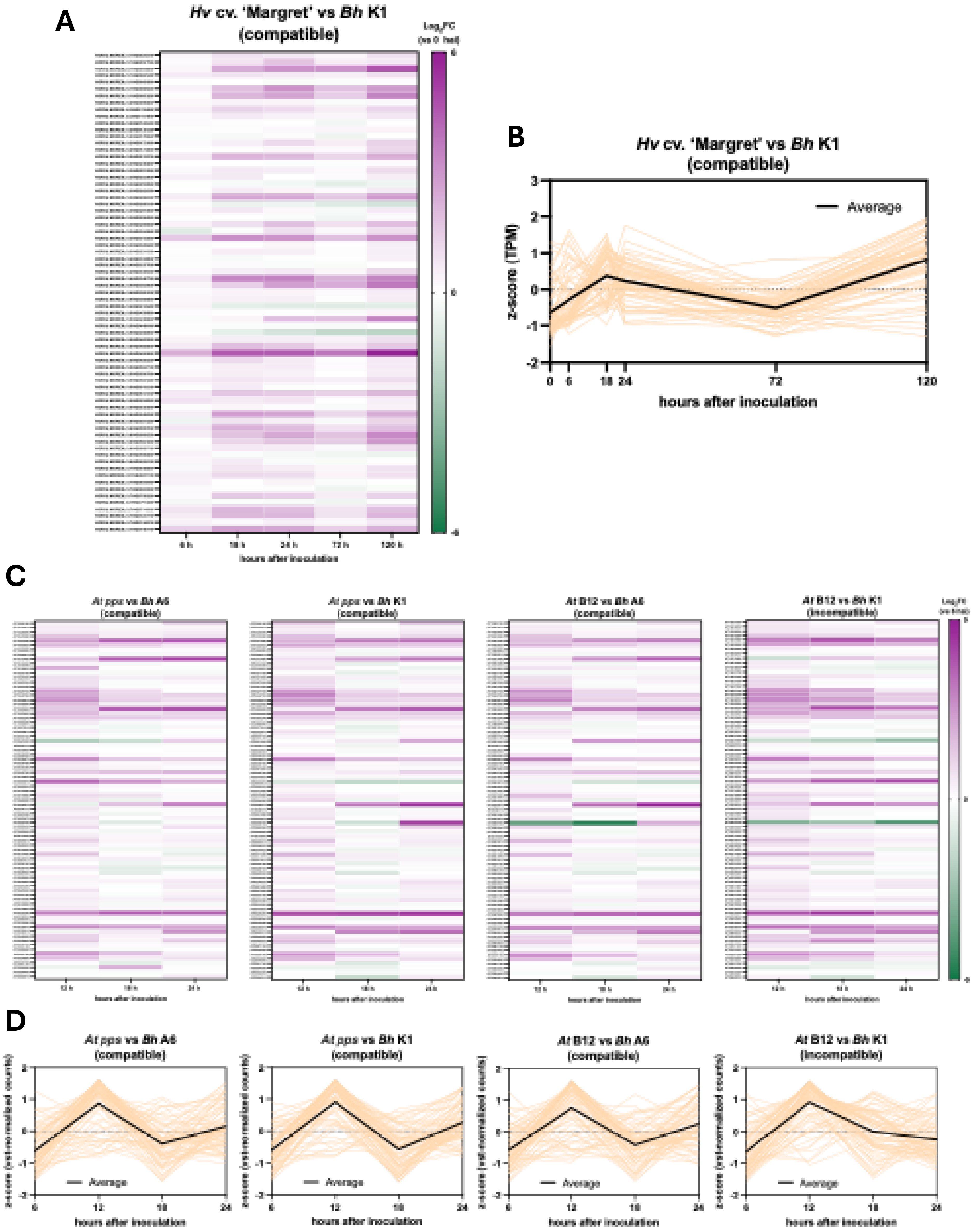
A
Hv cv. 'Margret' vs Bh K1
(compatible)
hours after inoculation
B
Hv cv. 'Margret' vs Bh K1
(compatible)
Average
z-score (TPM)
hours after inoculation
C
At pps vs Bh A6
(compatible)
At pps vs Bh K1
(compatible)
At B12 vs Bh A6
(compatible)
At B12 vs Bh K1
(incompatible)
D
At pps vs Bh A6
(compatible)
At pps vs Bh K1
(compatible)
At B12 vs Bh A6
(compatible)
At B12 vs Bh K1
(incompatible)
Average
hours after inoculation

**Supplemental Figure 11. The expression levels of genes commonly upregulated during ER stress peaks at the onset of haustoria formation in barley and partially immunocompromised Arabidopsis attacked by the *Bh* fungus.** Analysis of RNA-seq datasets of susceptible barley (*H. vulgare* cv. 'Margret') infected with the *Bh* isolate $K1_{AC}$ (**A and B**; Qian et al., 2023) and Arabidopsis *pen2-1 pad4-1 sag101-2* (*pps*) and B12 (*pps* 35S::MLA1-HA) infected with the *Bh* isolates K1 (containing $AVR_{A1}$) or A6 (lacking $AVR_{A1}$) (**C and D**; Maekawa et al., 2012). **(A)** Heat map showing the $\log_2$ fold change ($Log_2FC$) of individual genes commonly upregulated during the unfolded protein response (canonical UPR genes; Kim et al., 2018) at each timepoint relative to 0 hai. Each data point represents the mean of three independent repetitions. **(B)** Expression pattern of canonical UPR genes during *Bh* attack in barley. The z-score is based on transcripts per million (TPM). Black line represents the average expression of canonical UPR genes throughout the infection. Orange lines represent individual genes. **(C)** Heat maps showing the $Log_2FC$ of individual canonical UPR genes at each timepoint relative to 6 hai. **(D)** Expression pattern of canonical UPR genes during *Bh* attack in Arabidopsis. The z-score is based on vst-normalized counts. Black line represents the average expression of canonical UPR genes throughout the attack. Orange lines represent individual genes.

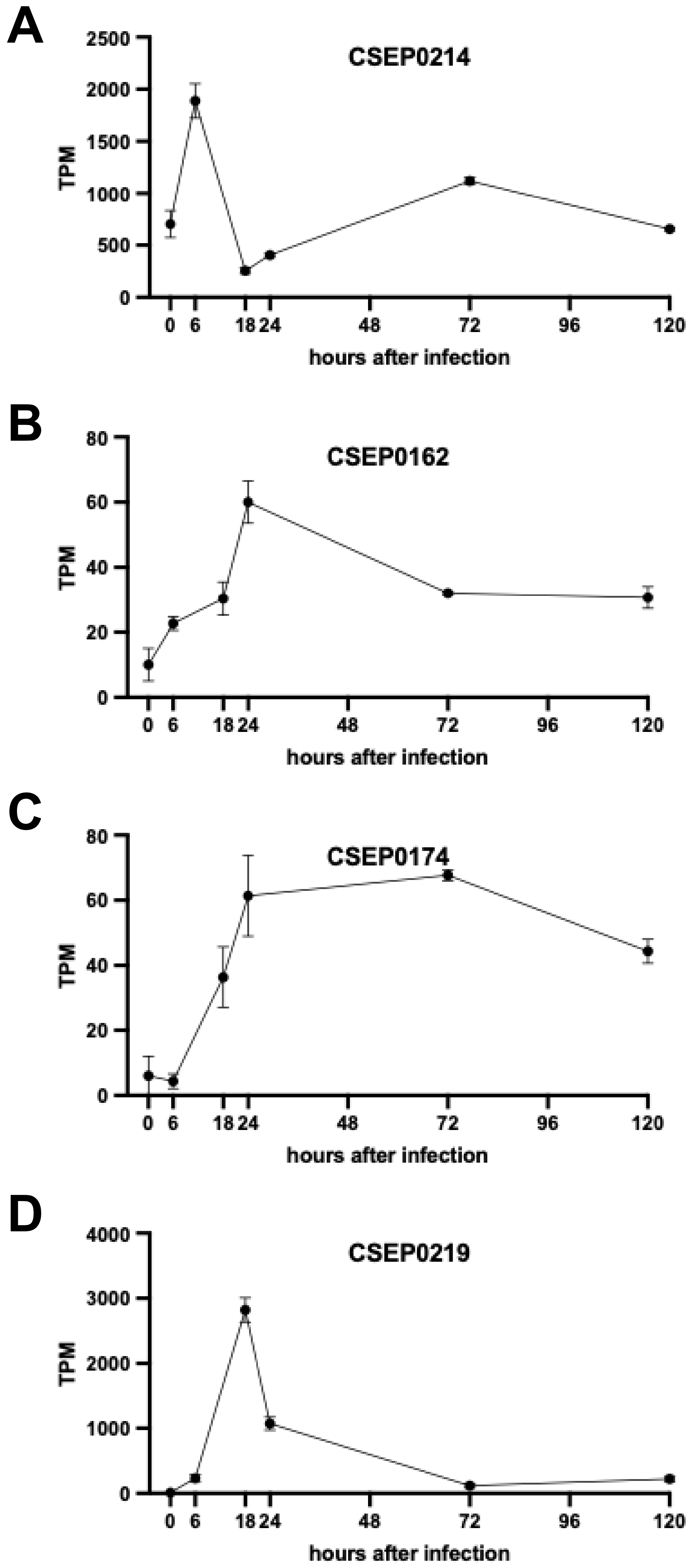


**Supplemental Figure 12. Expression patterns of vacuolar trafficking-suppressing CSEPs during barley infection by *Bh*.** The expression patterns of CSEP0214 **(A)**, CSEP0162 **(B)**, CSEP0174 **(C)**, and CSEP0219 **(D)** were retrieved from the RNA-seq datasets of susceptible barley (*H. vulgare* cv. 'Margret') infected with the *Bh* isolate $K1_{AC}$ (Qian et al., 2023). Expression is represented as transcripts per million (TPM). Each data point represents the mean of three independent repetitions. Error bars represent SD.

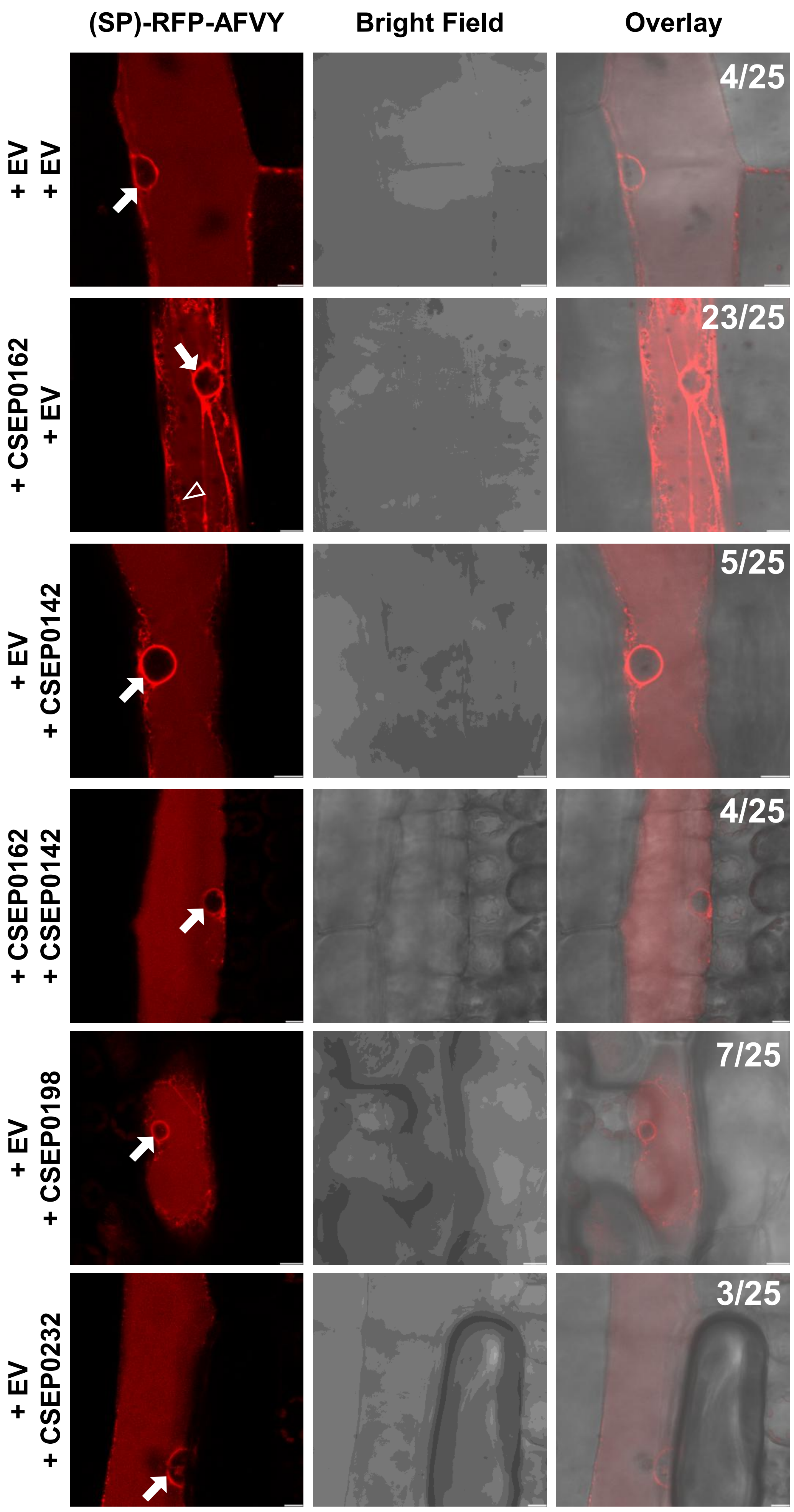


**Supplemental Figure 13. Release of barley powdery mildew effector CSEP0162-mediated ER-stalling by its interacting effector, CSEP0142.** Vacuolar marker, (SP)-RFP-AFVY, stalled in ER with reticular signal upon CSEP0162 expression. CSEP0142, CSEP0198 and CSEP0232 expressions cause minor (SP)-RFP-AFVY reticular signal in this particular experiment. Co-expression of CSEP0142 with CSEP0162 prevents the reticular signal. EV, empty vector. Numbers indicate proportions of epidermal cells showing major (SP)-RFP-AFVY reticular signal. Particle bombardment was carried out on 7-day-old barley *cv.* Golden Promise leaves and cells were imaged 2 days later. Open arrowhead, major reticular signal. Closed arrow, perinuclear signal. Scale bars, 10 µm. These results are representative of the outcomes of two independent experiments.
