## Supplementary figures and images for "Powdery mildew fungi block plant vacuolar traffic to suppress immunity"

### Source data Supplemental Fig. 6A

#### α – HA blot

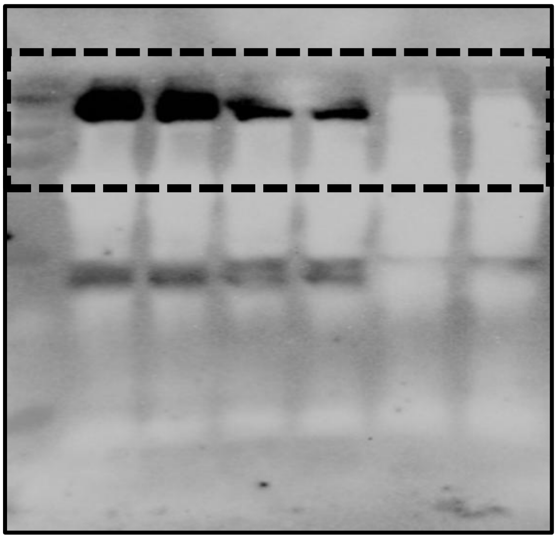

#### Ponceau staining for α – HA blot

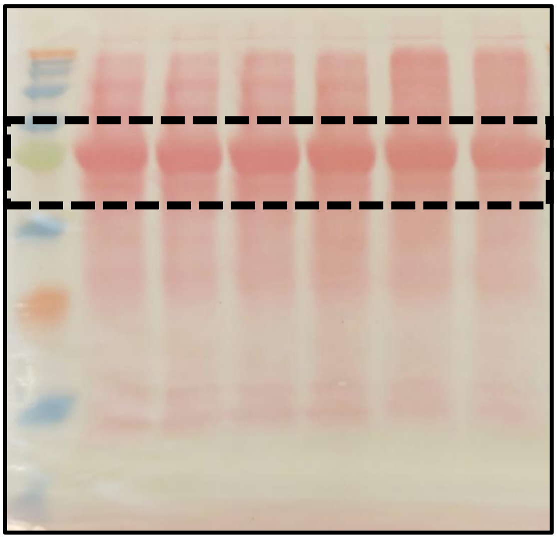

#### α – Flag blot

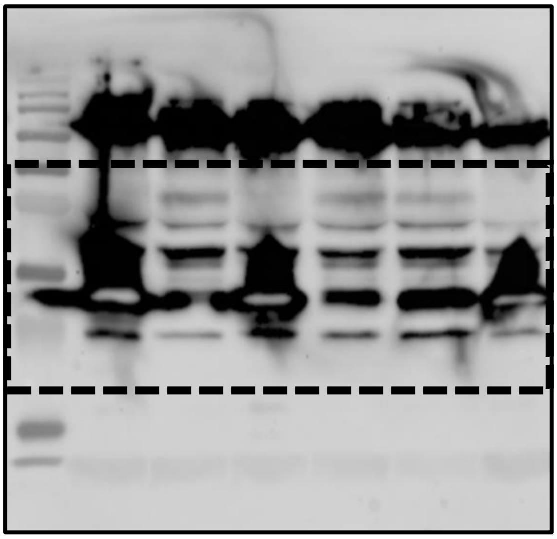

#### Ponceau staining for α – Flag blot

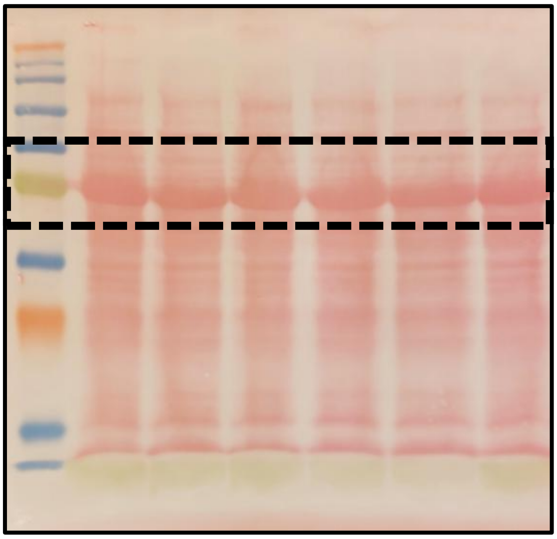
